# *In vivo* diversification of target genomic sites using processive T7 RNA polymerase-base deaminase fusions blocked by RNA-guided dCas9

**DOI:** 10.1101/850974

**Authors:** Beatriz Álvarez, Mario Mencía, Víctor de Lorenzo, Luis Ángel Fernández

## Abstract

Diversification of specific DNA segments typically involve *in vitro* generation of large sequence libraries and their introduction in cells for selection. Alternative *in vivo* mutagenesis systems on cells often show deleterious offsite mutations and restricted capabilities. To overcome these limitations, we have developed an *in vivo* platform to diversify specific DNA segments based on protein fusions between various base deaminases (BD) and the T7 RNA polymerase (T7RNAP) that recognizes a cognate promoter oriented towards the target sequence. The transcriptional elongation of these fusions generates transitions C to T or A to G on both DNA strands and in long DNA segments. To delimit the boundaries of the diversified DNA, the catalytically dead Cas9 (dCas9) is tethered with custom-designed crRNAs as a “roadblock” for BD-T7RNAP elongation. While the efficiency of this platform is demonstrated in *E. coli*, the system can be adapted to a variety of bacterial and eukaryotic hosts.

## Introduction

Directed evolution enables the selection of protein variants with improved properties as therapeutics and biocatalysts ^1, 2, 3^. The generation of genetic variability followed by a screening process are the essential steps of directed evolution ^4, 5^. *In vitro* mutagenesis techniques (e.g., error-prone PCR) can quickly produce large number of variants of the target gene but their selection requires, in most cases, cloning and transformation into a host cell for expression (e.g., *E. coli*). These steps are time-consuming and labor-intensive, especially when iterative cycles of mutagenesis and selection are needed. Cell-free selection methods are also feasible ^6^, but they are technically demanding and functional expression of complex proteins (e.g. membrane proteins, multimeric enzymes) is often difficult to achieve. Hence, *in vivo* mutagenesis systems are preferred because the generation of genetic variants, their expression and selection can done in a continuous process, which accellerates directed evolution ^5, 7^.

Long-established *in vivo* mutagenesis methods (e.g., chemical mutagens, radiations) do not target specific genes and are deleterious for the host cell due to accumulation of random mutations in the genome ^8^. Similarly, “mutator” host strains do not allow the concentration of the mutagenic activity on the target gene, inducing the accumulation of unwanted mutations in the host genome ^9, 10, 11, 12^. A few *in vivo* mutagenesis systems with targeted specificity have been reported for *E. coli* and yeast cells, which are the preferred hosts for cloning and expression of gene libraries ^5^. For example, an error-prone variant of *E. coli* DNA polymersase I enables the mutagenesis of cloned genes in ColE1 plasmids, although it concentrates the mutations close to the origin of replication ^13^. A different approach involves the transformation of *E. coli* with mutant oligonucleotide libraries, targeting one or multiple loci in the chromosome, which induce mutation during DNA replication *in vivo* ^14, 15, 16^. These systems allow multiloci genome engineering, but also imply iterative cycles of transformation with oligonucleotide libraries followed by high-throughput screenings, often including massive DNA sequencing steps, which are labor-intensive and demand sophisticated equipment. In yeast, generation variability can be achieved by cloning the target DNA segments in retrotransposons having an error-prone retrotranscriptase ^17^, but this process is limited to the DNA size that can be cloned in the retrotransposon (<5 kb).

A more versatile mutagenic system for different hosts is based on the tethering of base deaminases (BDs), such as cytosine deaminasaes (CDs) and adenosine deaminases (ADs), to a target DNA using catalytically dead Cas9 (dCas9) or analogous enzymes targeted with a CRISPR RNA (crRNA) or guide RNA (gRNA) ^18, 19, 20, 21^. Previous studies had shown that expression of CDs, such as human AID and orthologs from rat (rAPOBEC1) and lamprey (pmCDA1), induce random C to T mutations *in vivo*, both in *E. coli* and yeast. These CDs increase the frequency of C:G to T:A base pair transitions in DNA, especially when uracil DNA N-glycosilase (UNG) activity is inactived by gene deletion or by the specific inhibitor UGI ^22, 23, 24, 25^. This is because cytosine deamination produces uracil in DNA that can be eliminated by UNG, generating an abasic DNA that is a substrate of the base-excision repair system ^26, 27^. When a CD is fused to dCas9, its mutagenic activity is tethered to the target DNA sequence hybridized by the crRNA (or gRNA) allowing edition of specific bases in the genome ^18, 19, 20, 21^. In addition, mutations of A to G (inducing A:T to G:C base pairs transitions) have been generated by fusing dCas9 to an engineered AD named TadA*, derived from the endogenous RNA-dependent AD TadA of *E. coli* ^28^. Fusions of these BDs to dCas9 provides precise molecular tools for edition of specific bases in the genome, but its lack of processivity limits its potential for directed evolution of complete genes and operons. An interesenting alternative is the use of a nickase Cas9 (nCas9) fused to an error-prone DNA polymerase that is able to introduce mutations in a DNA segment of up to 350 bp ^29^, which is still limited for long genes and operons.

Hence, despite the above-mentioned advancements, the development of *in vivo* mutagenesis systems with high specificity and processivity are of great interest. In this study we report a novel approach that fulfill these criteria enabling *in vivo* mutagenesis of a target gene using different highly processive protein fusions of BDs and the bacteriophage T7 RNA polymerase (T7RNAP). The specificity of the mutagenesis is provided by a distinct T7RNAP promoter (P_T7_) driving transcription through the target DNA. By placing the P_T7_ at the 3’-end of the target gene, in reverse orientation, expression of the target gene can be preserved from its endogenous 5’-promoter recognized by the host RNAP. We report the mutagenic action of fusions of T7RNAP with different BDs (i.e., AID, rAPOBEC1, pmCDA1 and TadA*) on a target genomic DNA segment and show that the DNA-bound crRNA/dCas9 complex hinders elongation BD-T7RNAP hybrids, protecting the downstream DNA. Given the demonstrated functionality of BDs, T7RNAP and dCas9 in different hosts (e.g., bacteria, yeast, mammalian cells) ^18, 19, 21, 30, 31, 32^, this system has the potential to be implemented in diverse organisms other than *E. coli*.

## Results

### An E. coli reporter strain of the mutagenic and transcriptional activity of BD-T7RNAP fusions

A scheme of the overall strategy followed in this study is shown in Fig. 1. To measure both the mutagenic and transcriptional activity of BD-T7RNAP fusions, we designed a GFP-URA3 genetic cassette comprising two gene reporters in reverse orientation: a promoter-less *gfp* gene and the URA3 gene from *Saccharomyces cerevisiae* (Fig 2a). Transcription of the URA3 gene was placed under the Ptac promoter recognized by *E. coli* RNAP. Yeast URA3 encodes the enzyme orotidine 5’-phosphate decarboxylase involved in the synthesis of uridine monophosphate (UMP) ^33^. The activity of URA3, and that of the *E. coli* orthologue *pyrF*, allows cell growth in the absence of uracil in the medium (positive selection). In addition, URA3 expression makes yeast and *E. coli* cells sensitive to 5’-Fluororotic acid (FOA) allowing selection of null mutants (negative selection or counterselection) ^34, 35^. To enable specific recruitment of BD-T7RNAP fusions, the promoter P_T7_ was placed downstream of URA3, in reversed orientation to the coding sequence of URA3, but in the same orientation that the promoter-less *gfp* gene (Fig. 2a). Thus, expression of GFP acts as a reporter of the transcriptional activity of BD-T7RNAP fusions. The GFP-URA3 cassette was flanked by transcriptional terminators (T1 and T0). The whole genetic construct was cloned in an integrative suicide vector carrying an apramycin-resistance marker (Apra^R^) and flanking homology regions of the *flu* gene of *E. coli* K-12, encoding Antigen 43 ^36^. Integration of genetic constructs replacing *flu* does not affect bacterial growth and viability ^37, 38^. The *E. coli* K-12 strain used for integration was derived from the reference strain MG1655 ^39, 40^ by correcting using recombineering a natural mutation that reduces the expression of *pyrE* ^41^ (Supplementary Fig. 1a). The *pyrE* gene encodes the enzyme orotate phosphoribosyltransferase required for the biosynthesis of UMP and the incorporation of FOA to produce the toxic 5-FUMP. The strain with the corrected *pyrE* allele (named MG1655*) showed higher sensitivity to FOA than the parental MG1655 strain (Supplementary Fig 1b). Deletion of *pyrF* in MG1655* makes bacteria resistant to FOA (Supplementary Fig. 1b). The GFP-URA3 cassette was integrated in the chromosome of MG1655*Δ*pyrF* generating the reporter strain MG*-URA3, which grows well in mineral media (M9) lacking uracil and is highly sensitive to FOA (Supplementary Fig. 1b). Lastly, an *ung* deletion mutant was obtained in this strain (MG*-URA3Δ*ung*).

**Figure 1.**
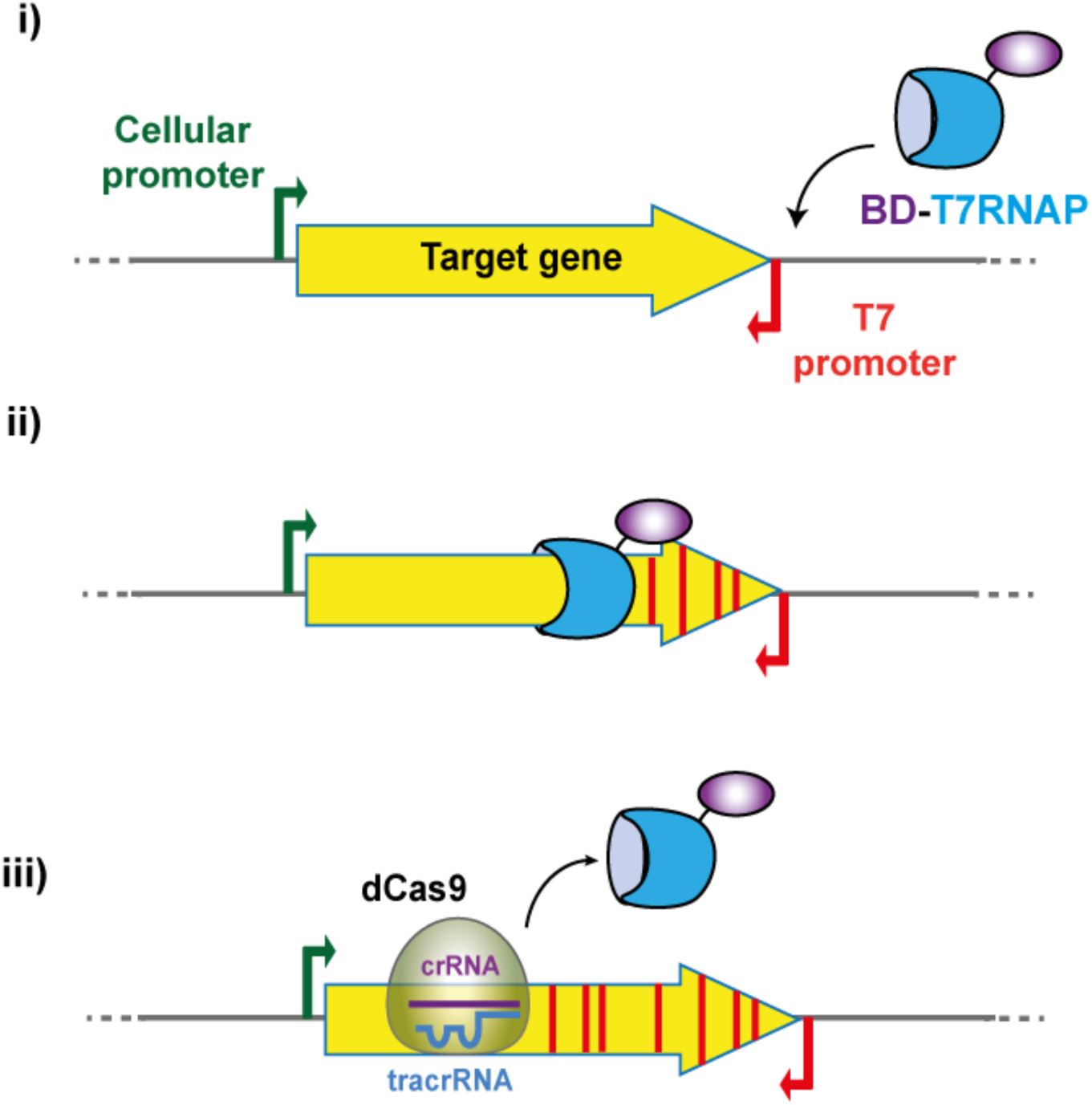
Schematic representation of the mutagenic process. The T7 RNA polymerase of the fusion specifically binds the T7 promoter (i), initiating the transcription and moving along the target gene carrying the base deaminase (BD) that creates mutations (red stripes) in this gene (ii). The enzymatic fusion stops and detaches from the DNA when encounters a dCas9 molecule bound to a specific sequence determined by the crRNA (iii).

**Figure 2.**
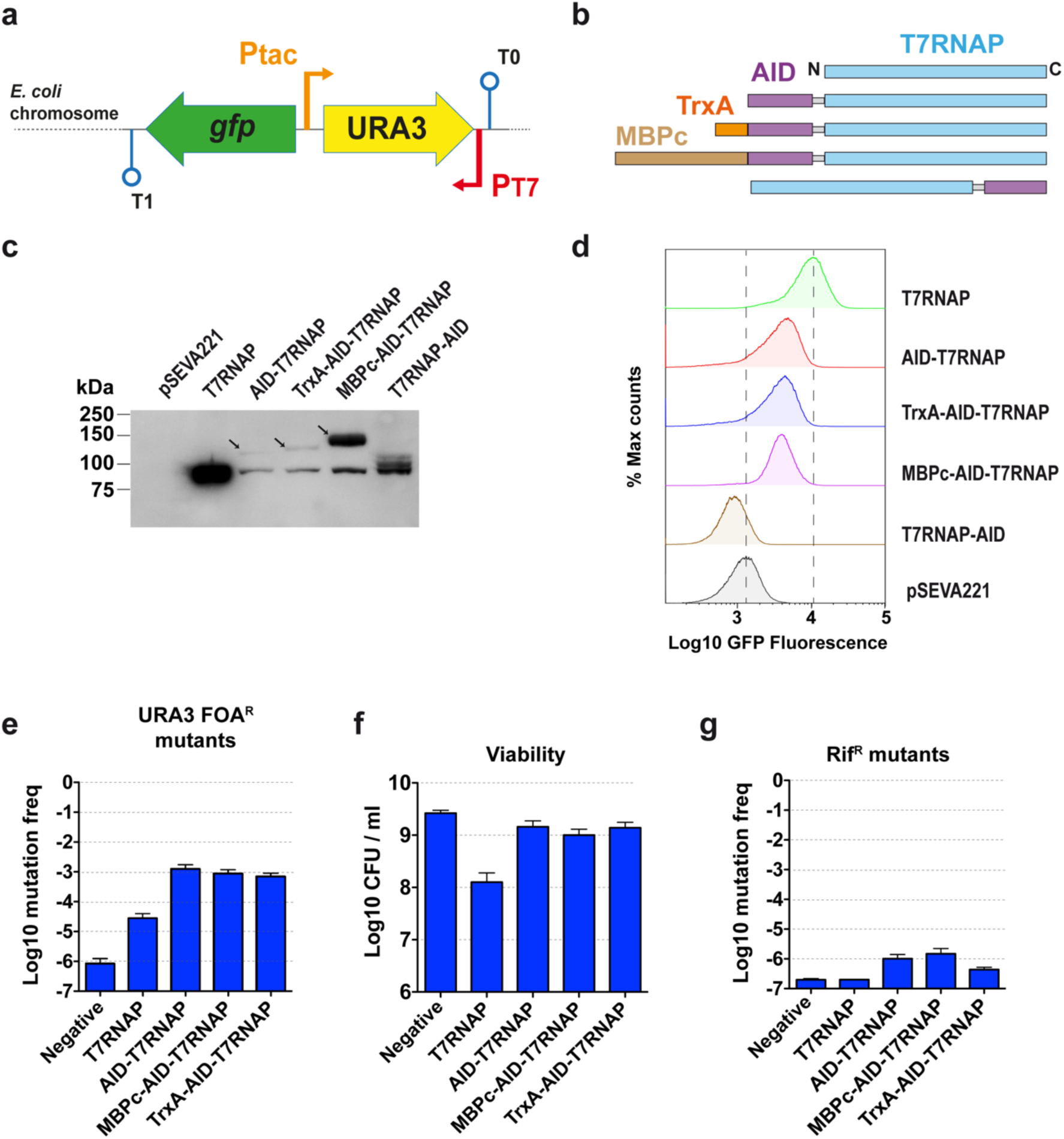
Expression and activity of the different variants of the fusion AID-T7RNAP. (**a**) Representation of the chromosomally integrated reporter cassette to test the mutagenesis system. Thin arrows indicate the promoters tac (P_tac_) and T7 (P_T7_), lollipops indicate terminators T0 and T1. (**b**) Representation of the different AID fusion variants. (**c**) Expression of the different variants determined by Western blot analysis of the cell extracts from induced cultures of the strain MG*-URA3Δ*ung* transformed with the different plasmids. The arrows indicate the bands corresponding to the full-length fusions. (**d**) Processivity of the fusions assessed by flow cytometry analysis to detect expression of *gfp* in the induced cultures. (**e**) Mutation frequency of URA3 as the ratio of FOA^R^ CFU/ml *vs*. total CFU/ml. (**f**) Viability as Log_10_ CFU/ml. (**g**) Mutation frequency of *rpoB* as the ratio of Rif^R^ CFU/ml *vs*. total CFU/ml. The histograms (**e, f** and **g**) represent the means and standard errors of three independent experiments (n=3).

### Expression and activity of T7RNAP fusions to human AID

We first investigated the tolerance of T7RNAP (∼99 kDa) for N- and C-terminal protein fusions using the human CD AID (∼24 kDa) ^22,24^. AID was fused at the N- and C-terminal ends of T7RNAP using a flexible peptide linker of Gly and Ser (G3S)7, generating AID-T7RNAP and T7RNAP-AID fusions, respectively (Fig. 2b). In addition, variants of AID-T7RNAP were constructed with different N-terminal tags: thioredoxin 1 (TrxA; ∼11 kDa) and a cytosolic version of the maltose binding protein (MBPc; ∼40 kDa), generating TrxA-AID-T7RNAP and MBPc-AID-T7RNAP, respectively (Fig. 2b). N-terminal TrxA and MBPc are reported to increase the solubility and expression level of protein fusions in *E. coli* ^42, 43, 44, 45^. Gene constructs encoding T7RNAP and AID fusions were placed under the control of the tetracycline-regulated promoter (TetR-PtetA) ^46, 47^ in a low copy-number plasmid (pSEVA221) ^48^. MG*-URA3Δ*ung* bacteria harboring these plasmids, and pSEVA221 as negative control, were grown at 37 ^o^C in LB for 2 h (OD600 ∼1.0), and induced with anhydrotetracycline (aTc) for additional 1 h. All cultures grew to a similar final optical density (OD600 ∼2.5), except bacteria expressing native T7RNAP (OD600 ∼1.2) suggesting some toxicity of T7RNAP expression. Whole-cell protein extracts from these cultures were analyzed by Western blot with a monoclonal antibody (mAb) anti-T7RNAP (Fig. 2c). Protein bands corresponding to the expected size of full-length fusions and T7RNAP were detected in bacteria expressing all N-terminal AID fusions, albeit higher levels were found with MBPc-AID-T7RNAP. In the case of the C-terminal fusion T7RNAP-AID, multiple protein bands were visible corresponding to T7RNAP and truncated polypeptides derived from the fusion, but none corresponding to the full-length polypeptide (Fig. 2c). GFP expression was detected by flow cytometry in bacteria with native T7RNAP and all N-terminal AID fusions, but not in bacteria expressing the C-terminal fusion of AID or carrying the empty vector (Fig. 2d). GFP fluorescence was ca. 2-fold higher in bacteria expressing native T7RNAP than in bacteria expressing any of the N-terminal AID fusions (Fig. 2d). Therefore, N-terminal fusions to T7RNAP produce a transcriptionally active polypeptide in *E. coli*, whereas C-terminal fusions are not stable and transcriptionally inactive.

To test the mutagenic activity of the transcriptionally active fusions, cultures of MG*-URA3Δ*ung* bacteria carrying plasmid constructs encoding none (empty vector control), AID-T7RNAP, TrxA-AID-T7RNAP, and MBPc-AID-T7RNAP, were grown and induced as above. After induction, bacteria were plated on M9+uracil and M9+uracil+FOA to determine colony forming units (CFU/ml) in each media. The mutation frequency of URA3 was determined for each bacterial strain in three independent experiments as the ratio of FOA^R^ CFU/ml *vs*. total CFU/ml (Fig. 2e). These data revealed a significant increase (∼1000-fold) in the frequency of FOA^R^ bacteria (URA3 mutants) in cultures expressing AID fusions (∼10^-3^) compared to negative control bacteria (∼10^-6^). Interestingly, expression of native T7RNAP alone increased ∼20-fold (∼2×10^-5^) the frequency of URA3 mutants, suggesting some mutagenic activity caused by high-level transcription of URA3 and/or by the toxicity of T7RNAP overexpression, which also caused a ∼10-fold reduction in the total CFU/ml (Fig. 2f). In contrast, bacterial cultures expressing AID fusions did not show any significant change in total CFU/ml, with values similar to the control strain (∼10^9^ CFU/ml) (Fig. 2f). Hence, the high-frequency of URA3 mutants found in bacteria with AID fusions strongly suggests a mutagenic activity of the fusion polypeptides.

To evaluate the specificity of the mutagenic AID-T7RNAP fusions, bacteria from the above cultures were also plated on rifampicin (Rif)-containing plates. Rif^R^ colonies of *E. coli* are known to contain mutations in *rpoB*, encoding the β-subunit of *E. coli* RNAP ^49, 50^. The frequency of mutation in *rpoB* was determined for each culture as the ratio of Rif^R^ CFU/ml vs. total CFU/ml (Fig. 2g). These data revealed a ∼5-fold increase in the frequency of *rpoB* mutants in bacteria expressing AID fusions (∼10^-6^) compared to the negative control or bacteria expressing native T7RNAP (∼2×10^-7^) (Fig. 2g). These data are in accordance with previous work showing that expression of AID slightly increases the mutagenesis of non-specific loci in *E. coli* Δ*ung* strains ^24^. However, this low non-specific mutagenic activity of AID-T7RNAP fusions is clearly insufficient to explain the ∼1000-fold increase in URA3 mutants, suggesting a strong specificity of AID-T7RNAP fusions for the mutagenesis of URA3.

### Mutagenic activity of different CDs fused to the N-terminus of T7RNAP

Since all fusions having AID at the N-terminal of T7RNAP showed similar transcriptional and mutagenic activities (Fig. 2), we chose the AID-T7RNAP fusion, lacking any additional protein partner, to continue our work. We constructed similar N-terminal fusions with other CDs, namely pmCDA1 and rAPOBEC1 (Fig. 3a). As for AID, pmCDA1 and rAPOBEC1 fusions were also cloned in pSEVA221 under the control of TetR-PtetA. MG*-URA3Δ*ung* bacteria carrying pSEVA221 (negative control) or plasmids encoding T7RNAP, AID-T7RNAP, pmCDA1-T7RNAP, and rAPOBEC1-T7RNAP, were induced with aTc. Western blot analysis of whole-cell protein extracts from these cultures revealed similar expression levels of the fusion proteins, and higher expression of native T7RNAP (Fig. 3b), as before. Flow cytometry analysis also showed similar levels of GFP expression in bacteria encoding AID, pmCDA1 and rAPOBEC1 fusions, with MFI values roughly half of those found in bacteria expressing native T7RNAP (Fig. 3c). Hence, as for AID-T7RNAP, fusions pmCDA1-T7RNAP and rAPOBEC1-T7RNAP were transcriptionally active in *E. coli*.

**Figure 3.**
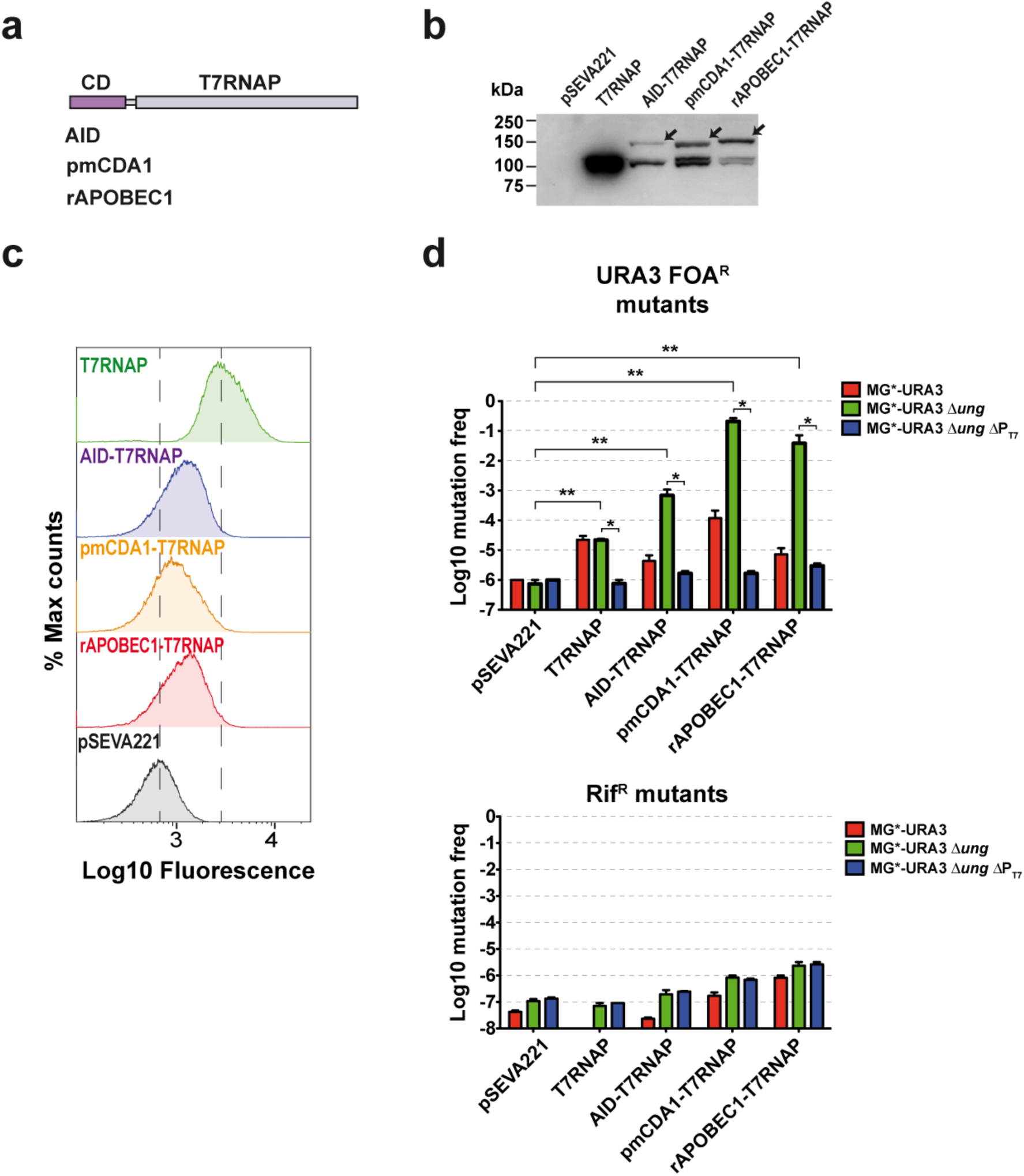
Expression and mutagenic activity of AID-T7RNAP, pmCDA1-T7RNAP and rAPOBEC1-T7RNAP. (**a**) Scheme of the different CDs fused to the T7 RNA polymerase by the linker (G_3_S)_7_. (**b**) Expression of the different fusions determined by Western blot analysis of the cell extracts from induced cultures of the strain MG*-URA3Δ*ung*. (**c**) Processivity of the fusions assessed by flow cytometry analysis to detect expression of GFP in the induced cultures. (**d**) Mutagenic activity in URA3 and *rpoB* of the different fusions using as hosts MG*-URA3, MG*-URA3Δ*ung* and MG*-URA3Δ*ung*ΔP_T7_. The histograms represent the means and standard errors of at least three independent experiments (n≥3). The statistical analysis was done using Mann Whitney test. Asterisks indicate p-value < 0.05 (*) and p-value < 0.01 (**).

Next, we determined the frequency of mutagenesis in URA3 and *rpoB* loci upon induction of bacterial cultures carrying pSEVA221 (empty vector) or derivatives encoding native T7RNAP, AID-, pmCDA1- and rAPOBEC1-fusions, in three different *E. coli* strains: MG*-URA3 (*ung*^+^), MG*-URA3Δ*ung*, and MG*-URA3Δ*ung*ΔP_T7_ (lacking the P_T7_ in URA3). The ratio of FOA^R^ CFU/ml and Rif^R^ CFU/ml *vs*. total CFU/ml for each induced culture was determined in three independent experiments (Fig. 3d). This analysis showed a significative increase in the mutagenesis of the URA3 locus for all CD fusions in Δ*ung* bacteria and in the presence of P_T7_. As expected from previous data (Fig. 2), the frequency of URA3 mutants in MG*-URA3Δ*ung* bacteria was ∼10^-3^ for AID-T7RNAP, ∼10^-5^ for native T7RNAP, and ∼10^-6^ for the empty vector, but increased dramatically to ∼5×10^-2^ for rAPOBEC1-T7RNAP and to ∼10^-1^ for pmCDA1-T7RNAP. Thus, rAPOBEC1 and pmCDA1 fusions have a higher mutagenic activity than AID fusion (∼50- to 100-fold, respectively). The higher activity of rAPOBEC1 and pmCDA1 was also partially reflected in a slighlty higher “off-target” mutagenesis in *rpoB* (Fig. 3d). Importantly, the frequency of URA3 mutants for all CD fusions dropped to levels close to those of the empty vector in the strain lacking the T7 promoter (MG*-URA3Δ*ung*ΔP_T7_ strain). Hence, these data clearly indicate that the strong mutagenic activity of CD-T7RNAP fusions in URA3 requires the presence of P_T7_. The requirement of P_T7_ is also seen for the mutagenic activity of native T7RNAP in URA3 (Fig. 3d). Lastly, we found that all CD fusions have a lower mutagenic activity in the *ung*^+^ strain whereas the mutagenic activity of native T7RNAP is independent of the presence of UNG (Fig. 3d). Not surprisingly, the highly active pmCDA1 fusion also showed the highest mutagenic frequency of URA3 in the *ung*^+^ strain, with a 100-fold increase compared to the control with the empty vector (frequency ∼10^-4^ vs. ∼10^-6^). A moderate increase of ∼5 to 10-fold in the frequency of URA3 mutants was found in the *ung*^+^ strain for AID and rAPOBEC1 fusions, respectively (Fig. 3d).

### Mutagenic activity of an adenosine deaminase (AD) fusion to T7RNAP

To broaden the capacity of the mutagenesis system to adenosine bases, a T7RNAP fusion was constructed with the modified AD TadA7.10 (TadA*) ^28^ (Fig. 4a). TadA* was shown to deaminate adenines in DNA generating inosines that leads to A:T > G:C transitions. The fusion TadA*-T7RNAP was expressed in MG*-URA3Δ*ung* at higher levels than AID-T7RNAP, as determined by Western-blot (Fig. 4b). The TadA*-T7RNAP fusion was also transcriptionally active, producing slightly higher levels of GFP than AID-T7RNAP (Fig. 4c). The potential mutagenic capacity of TadA*-T7RNAP was evaluated in different genetic backgrounds, using AID-T7RNAP as a positive mutagenesis control and pSEVA221 vector as a negative control (Fig. 4d). These experiments revealed that expression of TadA*-T7RNAP generates URA3 mutants with a frequency of ∼2-5×10^-4^ (∼100-fold higher than the negative control) in both MG*-URA3(*ung*^+^) and MG*-URA3Δ*ung* strains, indicating that TadA* fusion is mutagenic and, contrary to AID-T7RNAP, independent of the presence of the enzyme UNG (Fig. 4d). In *E. coli* K-12 the gene *nfi* encodes endonuclease V, which is involved in inosine elimination ^51, 52^. When TadA*-T7RNAP was expressed in *E. coli* strains lacking *nfi* (MG*-URA3Δ*nfi* and MG*-URA3Δ*ung*Δ*nfi*), the frequency of URA3 mutants further increased to ∼10^-3^, a level similar to that generated by AID-T7RNAP in the Δ*ung* mutant (Fig. 4d). In contrast, deletion of *nfi* had no effect over the mutagenesis frequency of AID-T7RNAP (Fig. 4d). In addition, expression of TadA*-T7RNAP did not produce any significant increase in the levels of off-target mutagenesis in *rpoB* in these strains compared to the negative control (pSEVA221) (Fig. 4d). Lastly, we confirmed that the mutagenic activity of TadA*-T7RNAP in URA3 was dependent on the presence of P_T7_ promoter since its mutagenesis frequency dropped to the baseline levels in the strain MG*-URA3Δ*ung*ΔP_T7_ (Fig. 4e). Altogether these data demonstrate that TadA*-T7RNAP fusion has a specific mutagenic activity for the target DNA having P_T7_. This mutagenic activity is independent of UNG, being moderately increased 2- to 5-fold when endonuclease V is absent.

**Figure 4.**
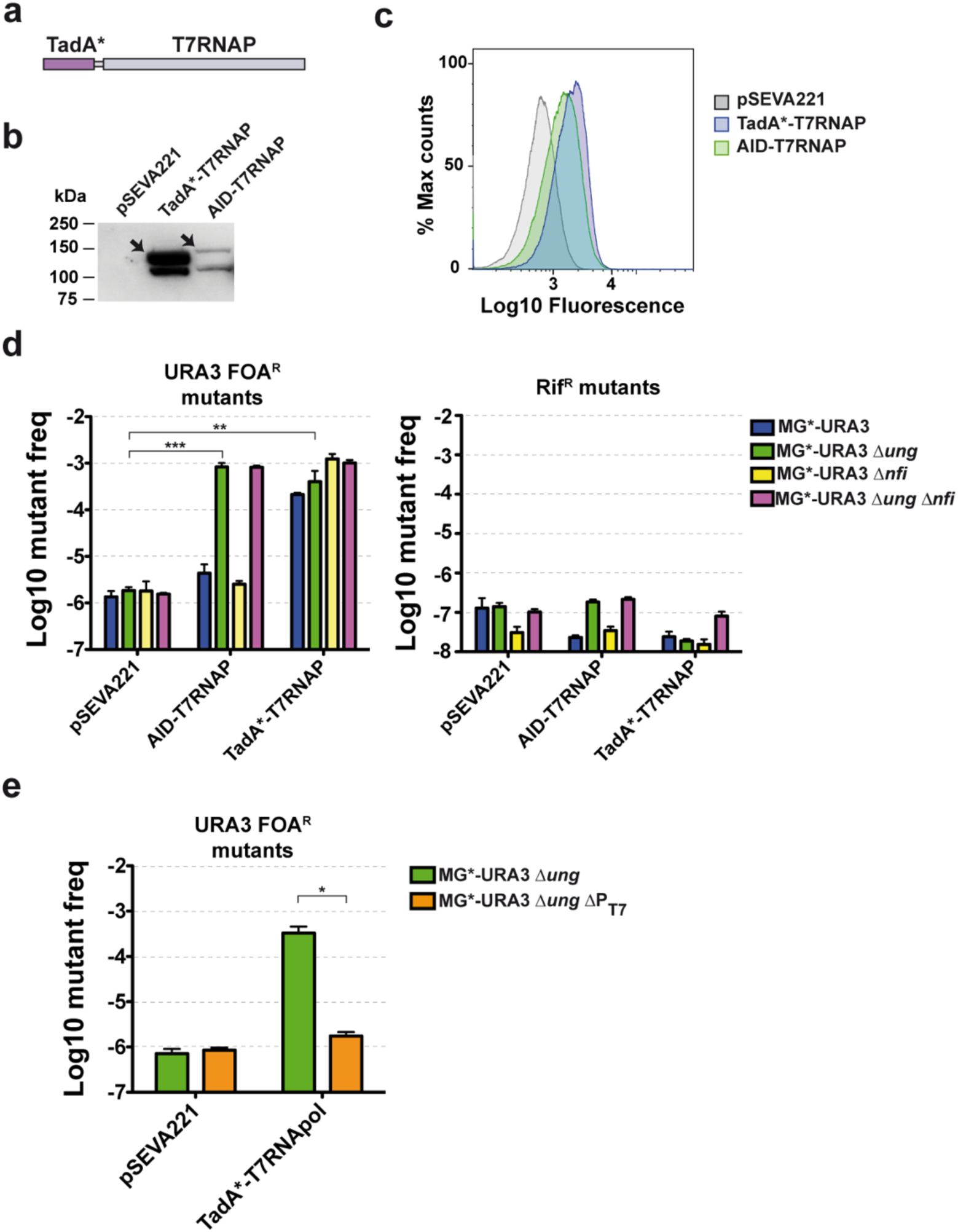
Expression and activity of TadA*-T7RNApol. (**a**) Scheme of the fusion TadA*-T7RNAP with the linker (G_3_S)_7_. (**b**) Expression of the fusion TadA*-T7RNAP in comparison to AID-T7RNAP determined by Western blot analysis of cell extracts from induced cultures. (**c**) Processivity of the fusions assessed by flow cytometry analysis to detect expression of GFP in the induced cultures. (**d**) Mutagenic activity of the AID-and TadA*-T7RNAP fusions in URA3 and *rpoB* using as hosts MG*-URA3, MG*-URA3Δ*ung*, MG*-URA3Δ*nfi* and MG*-URA3Δ*ung*Δ*nfi*. The histograms represent the means and standard errors of three independent experiments (n=3). (**e**) Mutagenesis frequency in URA3 when TadA*-T7RNAP is expressed in MG*-URA3Δ*ung* and MG*-URA3Δ*ung*ΔP_T7_ The histograms represent the means and standard errors of at least four independent experiments (n≥4). Asterisks indicate p-value < 0.05 (*), p-value < 0.01 (**) and p-value < 0.001 (***).

### Characterization of the mutations produced by BD-T7RNAP fusions

We randomly picked 30 FOA^R^ colonies (URA3 mutants) from the MG*-URA3Δ*ung* strains expressing native T7RNAP, AID-, pmCDA1- and rAPOBEC1-fusions to T7RNAP, and 30 additional FOA^R^ colonies grown from MG*-URA3Δ*ung*Δ*nfi* strain expressing TadA*-T7RNAP. The URA3 alleles from these colonies were amplified as PCR fragments including the 5’ Ptac promoter, the complete URA3 gene and downstream P_T7_, and their DNA sequence was determined. As an additional control, the URA3 alleles from 13 FOA-sensitive colonies from MG*-URA3Δ*ung* strain with plasmid pSEVA221 were amplified in parallel and sequenced, which did not show any mutation from the wild type sequence of URA3 allele. In contrast, in all FOA^R^ colonies analyzed from strains expressing CD-T7RNAP fusions we found multiple transitions C:G to T:A in both DNA strands along the Ptac and URA3 gene, but not in the P_T7_ sequence (Fig. 5a). In the case of FOA^R^ colonies from MG*-URA3Δ*ung*Δ*nfi* strain with TadA*-T7RNAP, all alleles contained transitions A:T to G:C in both DNA strands of the URA3 gene, except one different mutation, a C:G to T:A transition (Fig. 5b). No other type of DNA mutations, deletions, or insertions, were observed in any of the URA3 alleles of FOA^R^ colonies analyzed expressing BD-T7RNAP fusions. This was not the case in FOA^R^ colonies derived from the expression of native T7RNAP, which contained URA3 alleles with more types of mutations, including transitions (A:T to G:C and C:G to T:A), transversions (A:T to C:G and A:T to T:A), deletions, and insertions (Supplementary Fig. 2). Showing correlation to the mutagenic capacity of the three CD-T7RNAP fusions, the highest total number of mutations was found with pmCDA1-T7RNAP (426) followed by rAPOBEC1-T7RNAP (95) and AID-T7RNAP (42) (Fig 5c). The total number of mutations found in the 30 URA3 alleles from TadA*-T7RNAP was 37, similar to that of AID-T7RNAP (Fig. 5c). The average number of mutations per clone presented the same hierarchy: pmCDA1-T7RNAP (14.2) > rAPOBEC1-T7RNAP (3.2) > AID-T7RNAP (1.4) > TadA*-T7RNAP (1.2) (Fig. 5d). It is worth noting that for all CD-T7RNAP fusions, transitions G to A were detected more frequently than C to T in the URA3 coding strand (Figs. 5a and 5c). This indicates that CD-T7RNAP fusions mutates more frequently Cs in the non-coding strand of URA3, which corresponds to the non-template strand for the CD-T7RNAP fusions (Supplementary Fig. 3). This mutagenesis bias to the non-template strand is less pronounced for AID-T7RNAP (62%) than for rAPOBEC1-T7RNAP (74%) or pmCDA1-T7RNAP (91%) (Fig. 5c). In the case of TadA*-T7RNAP, we also found a bias favoring T to C mutations in the coding strand of URA3 (84%), corresponding to A to G mutations in the non-template strand for the TadA*-T7RNAP fusion (Fig. 5c and Supplementary Fig. 3). Therefore, DNA sequencing of URA3 mutant clones demonstrates that CD and AD fusions induce the expected mutations with a variable bias towards the non-template strand depending on the specific BD employed.

**Figure 5.**
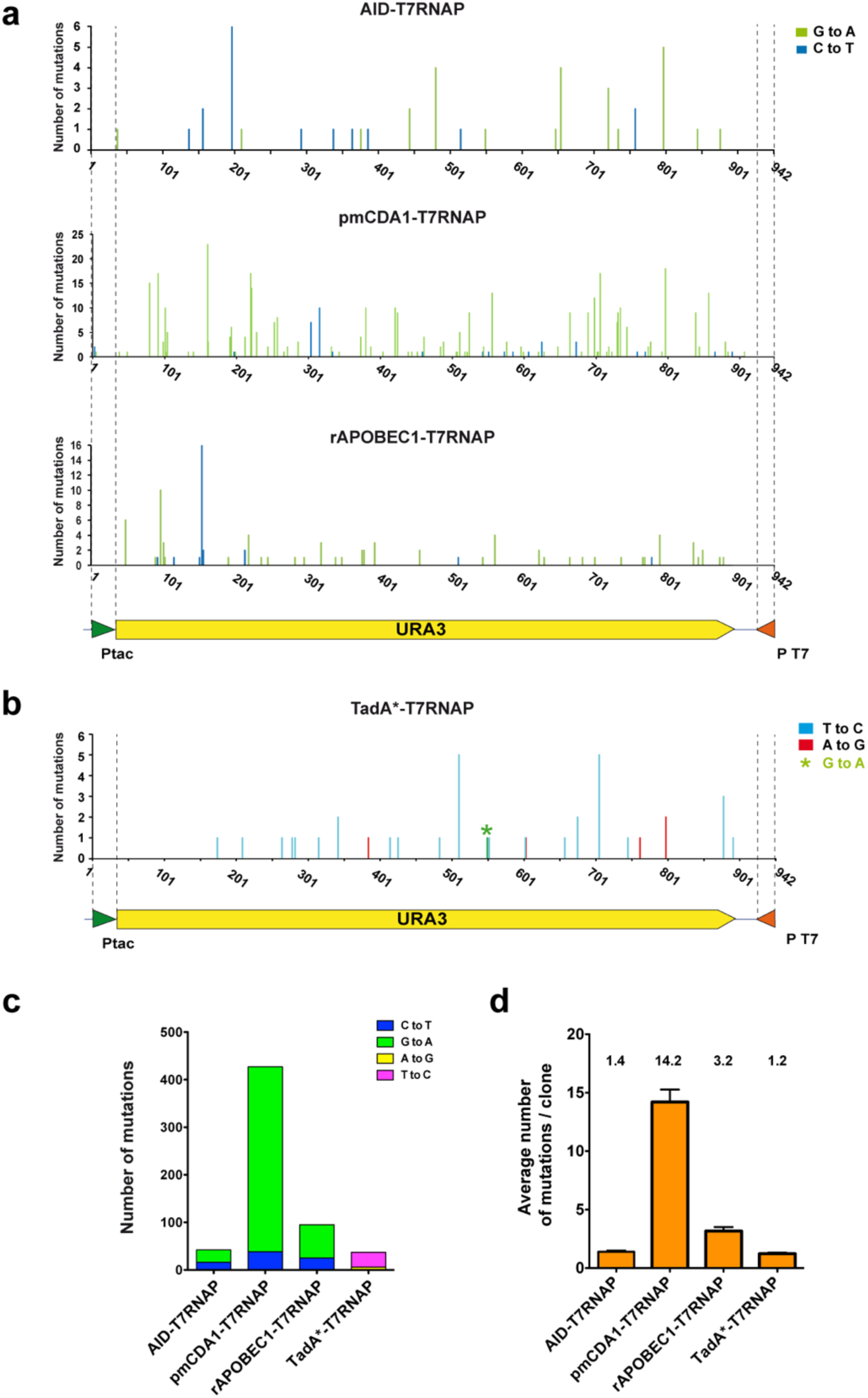
Characterization of URA3 mutations found in FOA^R^ colonies expressing BD-T7RNAP fusions. (**a**) Number of mutations per nucleotide identified in the URA3 locus from 30 FOA^R^-colonies isolated from each MG*-URA3Δ*ung* strain expressing the indicated CD-T7RNAP fusions and (**b**) from MG*-URA3Δ*ung*Δ*nfi* strain expressing TadA*-T7RNAP fusion. The promoters Ptac and T7 are shown with arrow heads and delimited by dashed lines. The indicated base changes correspond to the coding sequence of URA3. Different base substitutions found are labeled with the color codes on the right. A single G to A transition found with TadA*-T7RNAP is labelled with an asterisk. (**c**) Total number of mutations for each BD-T7RNAP fusion indicating the base substitutions found. (**d**) Average number of mutations per clone found in the FOA^R^-colonies analyzed for each of the indicated BD-T7RNAP fusions.

For a further analysis of the mutagenic process a 284 bp PCR fragment of the URA3 gene was amplified from a culture of MG*-URA3Δ*ung* expressing AID-T7RNAP and was subjected to massive next-generation sequencing (NGS; Materials and Methods). As a control, the same region was amplified from an induced culture of MG*-URA3Δ*ung* carrying the empty plasmid pSEVA221 and also subjected to NGS. The results of the variant call analysis from the two samples (ca. 1×10^6^ reads/sample) were compared and only the transition C:G to T:A appeared in an statistically significant higher number of reads in bacteria expressing AID-T7RNAP (Supplementary Fig. 4). Using an identical experimental approach, we analyzed the mutations caused in URA3 by TadA*-T7RNAP in MG*-URA3Δ*ung*Δ*nfi* compared with a culture of the same strain carrying pSEVA221. In this case, only transitions A:T to G:C were detected in a statistically significant higher number of reads in the sample expressing TadA*-T7RNAP (Supplementary Fig. 5). Hence, massive DNA sequencing data is consistent with the DNA sequencing results of individual URA3 mutants and confirms that the AID- and TadA*-T7RNAP generates only their expected transitions.

### Protection from mutagenesis of downstream regions by blockade with dCas9

The BD-T7RNAP fusions were able to transcribe the *gfp* gene downstream of URA3 and, presumably, creating mutations in this gene and beyond. To confirm this, we inserted in the GFP-URA3 cassette the *sacB* gene from *Bacillus subtilis* as an additional counter-selection system ^53^. The gene *sacB* codes for exoenzyme levansucrase, which uses sucrose to produce levan that accumulates in the periplasm of *E. coli* killing the bacteria. A Ptac-*sacB* fusion was inserted dowstream *gfp* in the GFP-URA3 cassette (Fig. 6a), and the new cassette was integrated in the *flu* locus of MG*Δ*pyrF*Δ*ung*Δ*nfi*. The resulting strain, named MG*-SacB-URA3Δ*ung*Δ*nfi*, was sensitive to sucrose with a frequency of spontaneous mutants of ca. 5×10^-6^. When AID-T7RNAP was expressed in this strain the mutagenesis frequency of *sacB* increased to ∼6×10^-4^ (Supplementary Fig. 6). The mutagenesis frequency of URA3 in this strain with AID-T7RNAP (∼1.7×10^-3^) was similar to that observed in MG*-URA3Δ*ung*Δ*nfi* (Fig. 4d). These results confirm that BD-T7RNAP fusions are able to mutate downstream regions to the target gene.

**Figure 6.**
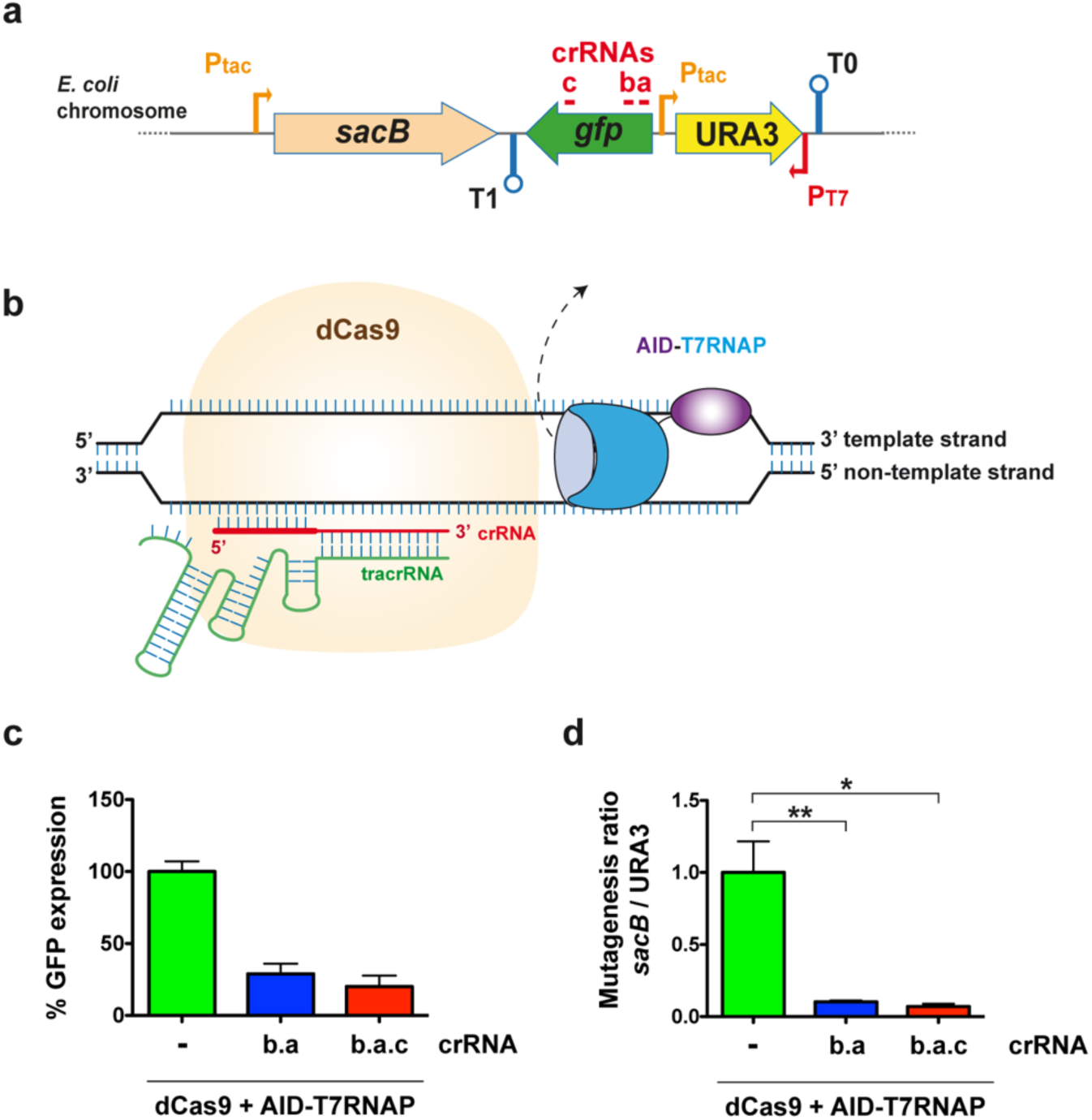
Blocking AID-T7RNAP elongation and mutagenic activity by dCas9 and crRNAs. **(a)** Scheme of the reporter cassette with *sacB* integrated in MG*-SacB-URA3Δ*ung*Δ*nfi* strain. Thin arrows indicate the tac (P_tac_) and T7 (P_T7_) promoters, lollipops indicate terminators T0 and T1, and red lines mark targeting sequences of the crRNAs a, b and c. **(b)** Representation of the dCas9 blocking activity showing one crRNA (in red) targeting the non-template strand relative to T7RNAP transcription. The mutagenic protein fusion is displaced from the transcription bubble (dashed arrow) by bound dCas9/crRNA. **(c)** Relative GFP levels measured by flow cytometry of bacteria from strain MG*-SacB-URA3Δ*ung*Δ*nfi* expressing AID-T7RNAP and dCas9 in the absence (-) or presence of crRNA arrays b.a and b.a.c. The histogram shows the percentages of mean flurorescence intensities (MFI) for each condition relative to the strain lacking crRNAs. Background GFP fluorescence signals from this strain with pdCas9 and the empty vector pSEVA221 are subtracted from all values. **(d)** Ratio of mutagenesis of *sacB* vs. URA3 in bacteria MG*-SacB-URA3Δ*ung*Δ*nfi* expressing AID-T7RNAP and dCas9 in the absence (-) or presence of crRNA arrays b.a and b.a.c. The ratio found in bacteria lacking crRNAs are considered 1. For (**c**) and (**d**), the histograms represent the relative means and standard errors from at least three independent experiments (n≥3). The statistical analysis was done using two-tailed Student t test. Asterisks indicate p-value < 0.05 (*), p-value < 0.01 (**).

To delimit the mutagenesis of BD-T7RNAP fusions within the target gene, we investigated the possibility of blocking elongation of T7RNAP with dCas9. dCas9 can bind a target DNA sequence using a crRNA or gRNA and has been successfully used as transcriptional repressor for the endogenous *E. coli* RNAP ^54^. Importantly, co-expression of two gRNAs targeting the same gene enhanced the transcription repression of *E. coli* RNAP by dCas9 ^55^. To study whether dCas9 can block elongation of BD-T7RNAP fusions we designed three crRNAs (Ta, Tb, Tc) against the non-template strand of *gfp* (Figs. 6a and 6b) and generate double (Tb.a) and triple (Tb.a.c) crRNAs. The plasmid pdCas9 ^54^ was used for the constitutive expression of dCas9, the transactivating CRISPR RNA (tracrRNA) and the designed crRNAs. The strain MG*-SacB-URA3Δ*ung*Δ*nfi* was transformed with plasmid pdCas9 (without crRNAs) and derivatives pdCas9-Tb.a and pdCas9-Tb.a.c. Then, these strains were transformed with the plasmid encoding AID-T7RNAP or the empty vector pSEVA221. After growth and aTc induction the levels of GFP expression were determined by flow cytometry (Fig. 6c). The expression of the double crRNAs Tb.a repressed GFP levels to ∼30% and with the triple crRNAs Tb.a.c the levels of GFP were further repressed to ∼20%. The basal level of GFP expression in strains carrying pSEVA221 was consider 0%. The protection of *sacB* from the BD-T7RNAP mutagenesis was assessed by the ratio of mutation frequencies in *sacB* vs. URA3 in each of these strains. This ratio was normalized as 1 for the strain expressing AID-T7RNAP and carrying pdCas9 without crRNAs (Fig. 6d). In concordance with the reduction of GFP expression detected by flow cytometry, the mutagenesis of *sacB* dropped ∼10 to ∼14-fold when the double (Tb.a) and triple (Tb.a.c) crRNAs were expressed, respectively (Fig. 6d). These results demonstrate that the dCas9 blockade can be used to limit the mutagenesis activity of BD-T7RNAP fusions mostly to a target gene (or gene segment), significantly reducing the mutagenesis of downstream DNA regions.

## Discussion

In this work we have reported an *in vivo* mutagenesis system with high specificity for a target gene based on the tethering of different BDs (AID, rAPOBEC1, pmCDA1 and TadA*) fused to T7RNAP, which selectively recognizes its specific promoter (P_T7_) in reverse orientation at the 3’-end and that transcribes thorughout the target gene. Expression of the target gene is maintained from its endogenous 5’-end promoter. The use of different BDs confers flexibility to the system due to their different mutagenesis profile. Although the expression level and transcriptional activity of CD-T7RNAP fusions were shown to be similar, their mutagenic capacity varied greatly ranging from the least active fusion with AID (URA3 mutagenesis frequency of ca. 10^-3^) to the most active fusion with pmCDA1 (URA3 mutagenesis frequency of ca. 10^-1^). The fusion bearing rAPOBEC1 produced an intermediate URA3 mutagenesis frequency of ca. 10^-2^. This means that the mutagenesis frequency of a particular target gene can be modulated using different CD-T7RNAP fusions. As expected, the mutations found in the target gene (URA3) by the CD-T7RNAP fusions were transitions C:G to T:A corresponding to the CD activity. In order to broad the mutation spectrum to A:T base pairs in the target gene, we also constructed the fusion TadA*-T7RNAP using the modified adenosine deaminase TadA7.10 (TadA*) ^28^. This fusion was expressed at higher levels than the CD-T7RNAP fusions, and was proved to generate bp transitions A:T to G:C in the target gene (URA3) with a mutagenesis frequency of ca. 10^-3^, similar to that elicited by AID-T7RNAP fusion. The average number of mutations in URA3 (ca. 1 kb) per clone agreed with the frequency of FOA^R^ mutants produced by the expression of the different fusions, being the fusion with pmCDA1 the enzyme introducing the highest number of mutations (ca. 14) whereas AID and TadA* fusions introduced the lowest number of mutations per clone (ca. 1.2 - 1.4). A single mutant isolated from TadA*-T7RNAP expression contained an unexpected bp transition (C:G to T:A), which could be caused by a non-specific base deaminase activity of TadA* or by the use of a common *E. coli* Δ*ung* host strain for expression of all BD-T7RNAP in these experiments. Nevertheless, massive DNA sequencing of a short amplicon of URA3, obtained after induction of AID and TadA* fusions without selection process, detected only the expected bp transitions C:G to T:A for AID and A:T to G:C for TadA*. Importantly, no other type of mutations (e.g. deletions, insertions) were found among the mutants isolated after expression of any of the BD-T7RNAP fusions. This result contrasts with the various types of mutations found after expression of T7RNAP alone. Our data revealed that transcription caused by T7RNAP alone increases the mutation frequency of the target gene by a mechanism that is independent of BDs, but that might be related to high exposure of ssDNA in the target gene and/or with conflicts of transcription with other cellular machineries (e.g. DNA polymerases during replication) ^56^.

Elimination of the DNA repair enzymes UNG and endonuclease V (*nfi*) was important to obtain optimal mutagenesis frequencies. It is well-known that UNG has a key role in reverting the mutations caused by CDs, so this enzymatic activity was inhibited in other mutagenesis systems based on CDs ^18, 19^. In our case, when UNG was present the mutagenesis frequency dropped between 500-fold for AID and 3000-fold for pmCDA1 and rAPOBEC. We deleted *ung* in our host strains to ensure complete abrogation of UNG activity, but an interesting alternative to *ung* deletion is its transient inhibition by expression of UGI (uracil N-glycosylase inhibitor) from bacteriophage PBS2 ^25^. For the mutagenesis with TadA*-T7RNAP, the effect of the endonuclease V was very mild. Only a decrease of 2- to 5-fold in the mutagenesis frequency with TadA*-T7RNAP was detected in bacteria with endonuclease V. Therefore, it seems that the removal of inosines generated by TadA* was much less efficient than removal of uracils generated by CDs in the genomic DNA of *E. coli*. This was also observed in eukaryotic cells subjected to mutagenesis with TadA* ^28^. It is worth to mention that UNG did not affect the TadA*-T7RNAP mutagenesis nor did endonuclease V affect the mutagenesis of AID-T7RNAP fusion.

Another observation to be noted is that the mutagenic capacity of the BD-T7RNAP fusions presented a bias towards the non-template strand. This phenomenon is less pronounced for AID-T7RNAP but it is detected with all of them. The reason of this strand preference is unclear but it may be caused by a better exposure of the cytosines or adenines on this strand in the transcription bubble. The template strand could be less accesible due its insertion in the catalytic core of the T7RNAP and the formation of the ssDNA:RNA hybrid ^57^.

A key property of a targeted mutagenesis system is the specificity towards the on-target sequence, keeping as low as possible the “off-target” mutagenesis to avoid the generation of deleterious mutations in other parts of the genome. The specifity of the system was assesed by two different approaches: monitoring mutations in the *rpoB* gene that conferred rifampicin resistance and monitoring mutations in the URA3 gene in absence of the T7 promoter. We found very high on-target vs. off-target ratios for all BD-T7RNAP fusions (ca. ≥ 10^3^). Nevertheless, some off-target mutagenesis is observed by the expression of these fusions. Compared to mutations frequencies found in a host strain (e.g. Δ*ung*) without BD-T7RNAP fusion, the “off-target” mutagenesis increased moderately (ca. 2 to 5-fold) for AID- and TadA*-fusions and more noticiable (ca. 10 to 20-fold) for pmCDA1- and rAPOBEC-fusions.

A different situation is caused by “off-target” mutations in downstream sequences (in relation to the T7 promoter) of the target gene due to the high processivity of the BD-T7RNAP fusions. To restrict the mutagenesis of BD-T7RNAP fusions to the target gene, we used dCas9 directed with crRNAs to block transcriptional elongation. This strategy has been previously used to repress gene expression in *E. coli* by blocking the endogenous RNA polymerase ^54, 55^. It has been reported that dCas9 blocks the *E. coli* RNAP when the crRNAs bind to the non-template strand and the presence of two targeting crRNAs enhance the repression ^55^. Keeping this in mind, we designed three crRNAs targeting the non-template strand. We have shown that the 3 crRNAs/dCas9 complexes are able to inhibit elongation of BD-T7RNAP fusions and reduce ca. 14-fold the mutagenesis of adjacent downstream DNA (with respect to T7RNAP elongation). Therefore, the protection from mutagenesis of dowstream regions was significant, yet it may be improved with dCas9 variants having increased blockade activity ^58^.

One interesting feature of the reported method is that it could be adapted for its use in different hosts since BDs, T7RNAP and dCas9 have been expressed in different bacteria, yeast and mammalian cells ^18, 19, 21, 30, 31, 32^. For modulation of the DNA repair systems, the expression of UGI would avoid the need of delete *ung* in host that are not *E. coli*. Elimination of *nfi* is not essential because the mutagenesis with TadA*-T7RNAP is still significant in the presence of the endonuclease V. However, the modified TadA* pairs with the endogenous TadA of *E. coli* to render a full active enzyme ^28^. Hence, for mutagenesis in hosts that lacks endogenous TadA, a heterodimeric construct fusing TadA and TadA* is needed ^28^. In mammalian cells incorporation of a nuclear localization signal (NLS) may be also necessary for targeting these protein fusions to the cell nucleus.

The system described in this work can be used for molecular evolution of different genes and operons of interest by simply replacing URA3 by these new target sequences. Alternatively the T7 promoter can be integrated downstream of genes or operons in the genome of *E. coli* or other cell host. The integration of multiple T7 promoters in different parts of the genome (e.g., downstream of genes involved in a metabolic pathway or in antibiotic resistance) would render in a multiplex genome editing for directed evolution of a cell phenotype. Alternatively, a plasmid that carries the target gene/s and the T7 promoter can be used. A recent independent study reported that a fusion between rAPOBEC1 and T7RNAP was able to generate mutations in antibiotic resistance genes located in plasmids carrying the T7 promoter ^59^. The use of multicopy plasmids can ease the initial utilization of the system but it has the drawback that creates multiple variants of the target gene/s in one cell. For selection of a particular variant, plasmid isolation and re-transformation of cells would be needed. When the genetic determinants are integrated in single copy in the genome, the mutagenesis and selection of the variants with the desired properties can be done in sequential iterative cycles without extensive manipulation of the culture as a continuos evolution process. We have demonstrate that different BDs-T7RNAP fusions can be used for the mutagenesis, and in particular that AID has lower off-target activity and mutagenic bias to the non-template strand than rAPOBEC1. Finally, our study also demonstrates that dCas9 with crRNAs can be used to limit the mutagenic activity within a target gene or gene segment without the introduction of multiple transcriptional terminators ^59^.

In sum, the BD-T7RNAP-based method hereby documented poses a simple workflow that could be applied for continuos molecular evolution of genes or operons with minimun manipulation. The extension of the mutagenized DNA can be delimited with dCas9 directed with designed crRNAs. Hence, *in vivo* mutagenesis with BD-T7RNAP fusions has the potential to be applied for the molecular evolution of biotechnologically relevant proteins, metabolic engineering of enzymatic pathways, diversification of gene libraries and other applications such as the study of the potential evolution pathways of antibiotic resistance genes.

## Methods

### Bacterial strains, media and growth conditions

A full list of the bacterial strains used in this study can be found in Supplementary Table 1. For plasmid propagation and cloning the *E. coli* strains used were DH10BT1R ^60^ and BW25141 (for the *pir*-dependent suicide plasmid derivatives) ^61^. All the strains were grown in Lysogeny Broth (LB) at 37 °C and shaking at 250 rpm ^62^. For solid medium, 1.5 % (w/v) agar was added to LB. The following antibiotics were added to the medium when needed: 50 μg/ml kanamycin (Km), 30 μg/ml chloramphenicol (Cm), 50 μg/ml apramycin (Apra) and 50 μg/ml rifampicin (Rif). Antibiotics were obtained from Duchefa-Biochemie. All other chemical reagents were obtained from Merck-Sigma unless indicated otherwise. For monitoring the mutagenic process, minimal medium M9 plates were used ^62^. This medium contains: 1x M9 salts (1 g/l NH_4_Cl, 3 g/l KH_2_PO_4_ and 6 g/l Na_2_HPO_4_), 2 mM MgSO_4_, 0.4 % (w/v) glucose, 0.0005 % (w/v) thiamine and 1.6 % (w/v) agar for solidification. When required, the minimal medium was supplemented with 20 μg/ml uracil and 250 μg/ml 5-fluorororic acid (FOA) (Zymo Research) or 60 g/l sucrose (counter-selection with *sacB*).

### Plasmids and cloning procedures

A list of the plasmids used in this study can be found in the Supplementary Table 2. The plasmids pGE*pyrF*, pGE*ung* and pGE*nfi* derived from *pir*-dependent plasmid pGE (Km^R^, R6K origin of replication) were constructed to delete the genes *pyrF, ung* and *nif*, respectively. Thermosensitive plasmids derivatives of pGETS (Km^r^, pSC101-ts origins of replication) were used to integrate the mutagenesis reporter cassettes in the non-essential locus of *flu*. The homology regions of the genes *pyrF, ung, nif* and *flu* were amplified by PCR using as template genomic DNA from *E. coli* MG1655. The gene *gfp*^*TCD* 63^ coding for GFP was obtained from the plasmid pGEyeeJPtac-gfp ^38^, the URA3 gene was a synthetic version optimized for *E. coli* (GeneCust), and the *sacB* gene was amplified from genomic DNA from *E. coli* T-SACK ^64^. The plasmid pSEVA221 (Km^r^, RK2-origin) ^65^ was used for expression of the T7RNAP fusions under the control of the *tetR*-P_tetA_ promoter ^46^. The *tetR*-P_tetA_ and the DNA fragments coding for the base deaminase (BD) enzymes fused to the linker peptide (G_3_S)_7_ were chemically synthesized with codon optimization for expression in *E. coli* (GeneART, ThermoFisher Scientific). The following BD enzymes were synthesized: human AID (Activation-induced cytidine deaminase), rAPOBEC1 (rat apolipoprotein B mRNA editing enzyme) ^66^, pmCDA1 (lamprey cytidine deaminase 1) ^67^, and TadA* (*E. coli* adenine deaminase variant TadA7.10) ^28^. The proofreading DNA polymerase Herculase II Fusion (Agilent Technologies) was used to amplify DNA fragments for cloning purposes. The plasmid pdCas9 ^54^ was used for the constitutively expression of the catalytically “dead” Cas9 (dCas9), the trans-activating crRNA RNA (tracrRNA) and the CRISPR RNAs (crRNA). The double (Tb.a) and triple (Tb.a.c) spacer arrays were cloned into the *Bsa*I site of pdCas9 using hybridized complimentary oligonucleotides, and following the one-step scheme CRATES ^68^ (Supplementary Methods). Cloning procedures were performed following standard protocols of DNA digestion with restriction enzymes and ligation ^62^. All DNA constructs were sequenced by the chain-termination Sanger method (Macrogen) and the sequences are deposited in GenBank (Supplementary Table 2).

### Generation of the reporter strains

The reporter strains for the mutagenesis system were derived from the reference of *E. coli* K-12 strain MG1655 ^39^. The strain MG1655* with higher sensitivity to FOA was generated by an oligo-mediated allelic replacement method ^16^ (Supplementary Methods). All the successive modifications were done over the MG1655* genetic background. The genes *pyrF, ung* and *nfi* were deleted using the plasmids pGE*pyrF*, pGE*ung* and pGE*nfi*, respectively, by a marker-less genome edition strategy ^37^. This strategy is based homologous recombination and resolution of the cointegrant promoted by the expression of the restriction enzyme I-*Sce*I and the λRed from the plasmid pACBSR ^69^. The URA3 cassettes were inserted in the *flu* locus using plasmid derivatives of pGETS, as reported previously ^38^.

### Induction of the mutagenesis system

A colony of freshly transformed reporter strain with the indicated plasmid derivative of pSEVA221, was grown overnight (O/N) in LB with Km at 37 °C with shaking (250 rpm). The next day, the culture was diluted 1:100 in fresh media and incubated under the same conditions for 2 h. Then anhydrotetracycline (aTc, 200 ng/ml) (TOKU-E) was added for induction and the cultures were incubated for 1 h. After that, a 500 μl-aliquot of each culture was washed with 1X phosphate-buffered saline (PBS) and resuspended in the same volume of PBS. A series of ten-fold dilutions of the cell suspension was prepared, and aliquots of 100 μl were plated in duplicates on different media: M9 + uracil; M9 + uracil and FOA; M9 + uracil and sucrose; and LB + Rif (see above for media composition).

### Flow cytometry analysis

Levels of expression of *gfp* from induced cultures were determined by flow cytometry analysis as follows: the volume corresponding to one unit of optical density (O.D.) at 600 nm of the induced cultures was collected by centrifugation (3300xg, 5 min) and resuspended in 500 μl 1X PBS. The cell suspension was diluted transferring 200 μl to a tube with 1200 μl of 1X PBS, and its fluorescence levels was determined using a Gallios FC500 flow cytometer (Beckman Coulter).

### Western blot analysis

To detect the expression of the fusion proteins in cell extracts of induced cultures, Western blot analysis was performed. From induced cultures, 0.5 O.D. was collected by centrifugation 3300 g 5 min, washed once with 500 μl 1X PBS, resuspended in 60 μl of PBS and mixed with 15 μl of 5X SDS-PAGE sample buffer ^70^. The samples were boiled 10 min before loaded into 8% polyacrylamide SDS gels, and electrophoresis was done using the Miniprotean III system (Bio-Rad) during 1 h 30 min at 170 V. The gels were then transferred to a polyvinylidene difluoride membrane (PVDF, Immobilon-P, Millipore) by means of O/N wet transfer (Bio-Rad) 4 °C at 30 V. The membranes were blocked with PBS 0.1% (v/v) Tween 20 with 3 % (w/v) skim milk powder and successively incubated with monoclonal mouse anti-T7 RNA polymerase antibodies (Novagen, Merck) and POD labelled goat anti-mouse antibodies (Sigma). Membranes were developed by chemiluminiscence using the Clarity Western ECL Substrate kit (Bio-Rad) and images were acquired using a ChemiDoc Touch system (Bio-Rad).

### DNA sequencing and analysis

To determine the DNA sequence of the URA3 alleles in the FOA resistant colonies, a DNA fragment of 1191 bp was amplified using the pair of primers F_GFPseq / R_T0ter (Supplementary Table S6) with the GoTaq Flexi DNA polymerase (Promega) by colony PCR following manufacturer’s instructions. The resulting amplicon was sequenced by the Sanger chain-termination method (Macrogen) using the same primers. The resulting 2 reads per colony were mapped against the Ptac-URA3-P_T7_ reference sequence (942 bp) to detect variants using the program SeqMan Pro (DNAstar).

For deeper analysis of the variations in URA3, NGS sequencing analysis of a 200 bp region was performed. To do this, genomic DNA was extracted from ∼4.5 ml of each induced culture (∼5 × 10^9^ bacteria) using the GNOME DNA Kit (MP Biomedicals). One hundred ng of total genomic DNA was used as template in a PCR reaction to amplify 284 bp of URA3 with the pair of primers F_CS1_URA3 / R_CS2_URA3 that includes Illumina tags CS1 and CS2 (Supplementary Table S6). The DNA amplification was carried out in 50 μl reactions using Herculase II Fusion DNA polymerase (Agilent Technologies) following manufacturer’s instructions. The amplicons were sent to the Genomic Unit of the Madrid Scientific Park to be sequenced by the NGS platform Illumina Miseq with paired-end (length > 2 × 300 bp) to acquire ∼1,000,000 reads per sample. These reads were processed with the program Bbmap (Bushnell et al., 2017, PMID: 29073143) for merging the paired-end reads. The resulting merged files were pilep up against the reference sequence using the program Samtools (Li et al., 2009, PMID: 19505943), and the variants were obtained with the program VarScan (Koboldt et al., 2012, PMID: 22300766) with the following parameters: --min-coverage 1 --min-reads2 1 --min-avg-qual 40 --min-var-freq 0.000001 --p-value 0.99. The sequences of the oligonucleotides used for amplification were discarded from the analysis since they may contain variations due to chemical synthesis.

### Statistics

Means and standard errors of experimental values were calculated using Prism 5.0 (GraphPad software Inc). Statistical analyses comparing groups in pairs were performed using Mann Whitney test (Fig. 3d, Supplementary Figs. S4 and S5) and two-tailed Student’s t-test (Figs. 4d and 4e, Fig. 6d) from at least three independent experiments. A value of p<0.05 was considered significant.

## Data availability

Data that support the findings of this work can be found in the main manuscript and in the Supplementary information. Materials and additional data are available from the corresponding author upon request. The sequences of the constructs built for this study are deposited in GenBank with the accession numbers listed in Supplementary Table 2.

## Acknowledgements

We thank Dr. David Bickard (Institute Pasteur, France) and Drs. Esteban Martínez, Yamal Al-ramahi, and Tomás Aparicio (CNB-CSIC) for providing some materials used in this work. The excellent technical work in massive DNA sequencing of the Genomic Unit of “*Parque Científico de Madrid*” is greatly appreciated. We are grateful to Dr. Alejandra Bernardini (Hospital 12 de Octubre Madrid, Spain) for assistance with massive DNA sequencing data analysis. This work is supported by the Grants BIO2017-89081-R (Agencia Española de Investigación AEI/MICIU/FEDER, EU) to LAF and ERC-2012-ADG_20120314 (European Research Council,EU) to VdL.

## Author contributions

LAF, MM and VdL conceived the study. BA and LAF designed the experiments and analysed the results. BA performed the experiments. All authors interpreted the data. BA and LAF wrote the initial manuscript and prepared figures. All the authors revised and approved the final manuscript.

