## Supplementary Information and Figures for "*In vivo* diversification of target genomic sites using processive T7 RNA polymerase-base deaminase fusions blocked by RNA-guided dCas9"

*(2) Centro de Biología Molecular "Severo Ochoa" (Consejo Superior de Investigaciones Científicas – Universidad Autónoma de Madrid), Nicolas Cabrera 1, Campus UAM Cantoblanco, 28049 Madrid, Spain*

---

* **Corresponding author**: Dr. Luis Ángel Fernández


### Supplementary Methods

#### Construction of the pdCas9 derivative plasmids pdCas9b.a and pdCas9b.a.c

For cloning the double Tb.a and the triple Tb.a.c spacer arrays into pdCas9 to generate the plasmids pdCas9b.a and pdCas9b.a.c, respectively, dsDNA fragments containing spacers and direct repeats (DRs) were built using hybridized complimentary oligonucleotides (Supplementary Fig. 7) that are listed in Supplementary Table 3. The sequences of the spacers used can be found in Supplementary Table 4. To generate pdCas9b.a, a dsDNA fragment containing spacer Tb-DR-spacer Ta with *Bsa*I cohesive ends was generated by annealing the pair of oligonucleotides 1 / 2 (Supplementary Fig. 8). To facilitate the cloning procedure, both oligonucleotides were purchased with phosphorylated 5' ends (Merck KGaA). To anneal the oligonucleotides, a mix was prepared in a PCR tube with 1 μl of 100 μM each oligonucleotide (100 pmoles), 2.5 μl of 1 M NaCl and 43 μl of water (Milli-Q). The mix was incubated in a thermocycler with the following annealing program: 1 cycle of 5 min at 95 ºC and 72 cycles of 1 min starting at 95 ºC and decreasing 1 ºC each cycle. The resulting dsDNA solution was kept on ice and diluted 1:10 in Milli-Q water to be used in a ligation mixture that contains 1 μl dsDNA diluted solution (0.4 pmoles), 2 μl pdCas9 digested with *Bsa*I (100 ng), 1 μl (1 U) T4 DNA ligase (Roche Diagnostics GmbH), 2 μl 10X T4 DNA ligase buffer and 14 μl of water (Milli-Q). The mixture was incubated O/N at 8 ºC, and subsequently used to electroporate *E. coli* DH10BT1R. From two resulting $Cm^R$ colonies, plasmid was isolated and the correct cloning of the double spacer array was confirmed by Sanger chain-termination method (Macrogen). The protocol for cloning the triple spacer array into pdCas9 was based on the one-step scheme CRATES (CRISPR Assembly through trimmed end of Spacers) described by Liao *et al.* [1]. In this case, three dsDNA fragments were assembled using the junction sequences GCTG and GAGT (Supplementary Fig. 8). These junction sequences were introduced replacing the first 4 nucleotides at 5' end of the spacers Ta and Tc. The new spacers with the junction sequences were named Ta* and Tc*. This modification does not affect the target specificity, since in the maturation process of the crRNAs, the spacers are trimmed approximately 10 nucleotides at the 5' end [2]. The three dsDNA fragments were: spacer Tb-DR (with annealed oligonucleotides 3 / 4), spacer Ta*-DR (with annealed oligonucleotides 5 / 6), and spacer Tc* (with annealed oligonucleotides 7 / 8) (Supplementary Fig. 8). Each oligonucleotide was 5' phosphorylated as follows: a mix was prepared with 1 μl of 100 μM oligonucleotide (100 pmoles), 5 μl of 10X T4 DNA ligase buffer (Roche Diagnostics GmbH), 1 μl (10 U) of T4 polynucleotide kinase (New England Biolabs) and water (Milli-Q) up to a total volume of 50 μl. The mix was incubated at 37 ºC 30 min, and then, the T4 polynucleotide kinase was inactivated at 65 ºC 20 min. The annealing process was carried out as described above, and 2 μl of each dsDNA fragment (0.2 pmoles) were added to a ligation mix containing 2 μl pdCas9 digested with *Bsa*I (100 ng), 1 μl (1 U) T4 DNA ligase (Roche Diagnostics GmbH), 2 μl 10X T4 DNA ligase buffer and 9 μl of water (Milli-Q). After incubation O/N at 8 ºC, the mix was used to electroporate *E. coli* DH10BT1R cells following standard procedures. From the resulting $Cm^R$ colonies, plasmid was isolated and the correct cloning of the triple spacer array was confirmed by DNA sequencing (Macrogen).

**Oligo-mediated allelic replacement method for the generation of MG1655***

The strain MG1655 was transformed with the plasmid pORTMAGE3 (Supplementary Table 2) that harbors the genes *gam*, *bet* and *exo* (λRed recombinase enzymes genes), and the *mutL E32K* (negative dominant variant of *mutL*). Expression of these genes is controlled by the cI857 temperature-sensitive repressor. An O/N culture of MG1655 with pORTMAGE3 was diluted 1:100 in 10 ml LB and grown at 30 ºC until mid-exponential phase (OD600 0.55-0.65), and then incubated at 42 ºC in a shaking water bath to induce λRed proteins and MutL (E32K) expression for 15 min 250 rpm. The induced culture was placed immediately on ice for at least 5 min, and the cells were pelleted and washed twice with 10 ml ice-cold Milli-Q water. The cells were resuspended in 160 μl of Milli-Q water and kept on ice. Forty microliters of cell suspension were electroporated with 1 μl of 100 μM rph oligo (Supplementary Table 5 and Supplementary Fig 9). As a control, another 40 μl cell suspension was electroporated with 1 μl TE (10 mM Tris-HCl, 1 mM EDTA, pH 8). Immediately after electroporation, 1 ml of SOC medium (20 g/l Bacto-tryptone, 5 g/l yeast extract, 0.58 g/l NaCl, 0.18 g/l KCl, 2 g/l MgCl2 and 1.2 g/l MgSO4) at room temperature was added and the cell suspensions were transferred to a flask containing 4 ml of SOC. Cells were allowed to recover for 60 min at 30 ºC 250 rpm, after that, 5 ml of LB was added and second cycle of electroporation was performed repeating the previous steps. After the second cycle of oligo recombineering, cells were collected by centrifugation, resuspended in M9 minimal medium and incubated O/N at 30 ºC 250 rpm. The next day, the cultures were diluted 1:$10^6$ and plated on minimal medium M9 lacking uracil for selection of clones harboring the insertion, since it was described that the modification affected bacterial growth in this medium [3]. After 24 h incubation at 30 ºC, a few larger colonies were observed in the sample that had been electroporated with rph oligo. Only small colonies were observed in the control sample. Twenty-three large colonies were tested for the presence of the extra G base with an allelic-specific PCR using the pairs of primers F_rph_A/R_rph (amplify 200 bp of *rph* allele w/o G) and F_rph_B/R_rph (amplify 200 bp of *rph* allele with G) (Supplementary Table 5). The allelic-specific PCR was carried out using GoTaq Flexi DNA polymerase (Promega) following the manufacturer's instructions. The optimized PCR program in the thermocycler was 1 cycle at 95 ºC for 5 min; 30 cycles at 95 ºC 30 s, 68.4 ºC 30 s and 72 ºC 30 s; and a final cycle at 72 ºC for 7 min. DNA samples from MG1655 and DH10BT1R were used as controls for DNA w/o the extra G and DNA with the extra G, respectively. Large colonies were tested by this allelic PCR and most of them produced an amplicon for the expected G insertion. This insertion was confirmed in two colonies by PCR amplification with the pair of primers F_seq_pyrE/R_rph, and subsequently by Sanger DNA sequencing (Macrogen) of the amplicons. The modified MG1655 strain was named MG1655*. The sensitivity of MG1655* to 5-FOA in comparison to the parental strain MG1655 was tested plating drops of ten-fold dilutions in 1 X PBS from liquid cultures on minimal medium M9 supplemented with uracil and 5-FOA.

#### Supplementary Table 1. *E. coli* strains used in this study

| Strain | Genotype and relevant features | Reference |
|---|---|---|
| **DH10B-T1R** | (F- λ-) *mcrA* Δ*mrr-hsdRMS-mcrBC* *φ80lacZDM15* Δ*lacX74* *recA1* *endA1* *araD139* Δ(*ara, leu*)7697 *galU* *galK* *rpsL* (StrR) *nupG* *tonA* | Novagen |
| **BW25141** | (F- λ-) Δ*(araD-araB)567*, Δ*lacZ4787*(::rrnB-3), Δ*(phoB-phoR)580*, *galU95*, Δ*uidA3*::*pir*, *recA1*, *endA9*(del-ins)*::FRT*, *rph-1*, Δ*(rhaD-rhaB)568*, *hsdR51* | [4] |
| **MG1655** | K-12 (F- λ-) | [5] |
| **MG1655*** | MG1655 *rph* (U00096:3815879_3815880insC)[a] | This work |
| **MG1655*Δ*pyrF*** | MG1655* Δ*pyrF* | This work |
| **MG*-URA3** | MG1655* Δ*pyrF* Δ*flu*::*gfp*-$P_{tac}$-URA3-$P_{T7}$-*aac(3)IV* | This work |
| **MG*-URA3Δ*ung*** | MG*-URA3 Δ*ung* | This work |
| **MG*-URA3Δ*nfi*** | MG*-URA3 Δ*nfi* | This work |
| **MG*-URA3Δ*ung*Δ$P_{T7}$** | MG1655* Δ*pyrF* Δ*ung* Δ*flu*::*gfp*-$P_{tac}$-URA3-*aac(3)IV* | This work |
| **MG*-URA3Δ*ung*Δ*nfi*** | MG*-URA3 Δ*ung* Δ*nfi* | This work |
| **MG*-SacB-URA3Δ*ung*Δ*nfi*** | MG1655* Δ*pyrF* Δ*ung* Δ*nfi* Δ*flu*::$P_{tac}$-*sacB*-*gfp*-$P_{tac}$-URA3-$P_{T7}$-*aac(3)IV* | This work |

Note: (a) Insertion of C between nucleotides 3815879 and 3815880 of genome accession number U00096

#### Supplementary Table 2. Plasmids used in this study

| Plasmid | Relevant features | Reference | Accession number |
|---|---|---|---|
| pORTMAGE3 | $Km^R$; pBRRI ori, *cI857*(ts), $P_R$ *exo*, *bet*, *gam*, *mutL E32K* | 6 | Not deposited |
| pACBSR | $Cm^R$; p15A ori, $P_{BAD}$, I-SceI endonuclease and λRed genes | 7 | Not deposited |
| pGE | $Km^R$; R6K ori, muticloning site and I-SceI sites | 8 | Not deposited |
| pGE*pyrF* | $Km^R$; ca.500 bp homolgy regions flanking *pyrF* cloned in pGE; for deletion of *pyrF* | This work | MN450165 |
| pGE*ung* | $Km^R$; ca.500 bp homolgy regions flanking *ung* cloned in pGE | This work | MN450166 |
| pGE*nfi* | $Km^R$; ca.500 bp homolgy regions flanking *nfi* cloned in pGE | This work | MN450167 |
| pGETS | $Km^R$; pSC101-ts ori, multicloning site and I-SceI sites | 9 | Not deposited |
| pGETS*flu*URA3 | $Km^R$ $Apra^R$; pGETS, *flu* HRs, *gfp*$^{TCD}$-$P_{tac}$-URA3-$P_{T7}$-*aac(3)IV* | This work | MN450168 |
| pGETS*flu*URA3($P_{T7}$)del | $Km^R$ $Apra^R$; pGETS, *flu* HRs, *gfp*$^{TCD}$-$P_{tac}$-URA3-*aac(3)IV* | This work | MN450169 |
| pGETS*flu*SacB-URA3 | $Km^R$ $Apra^R$; pGETS, *flu* HRs, $P_{tac}$-*sacB*-*gfp*$^{TCD}$-$P_{tac}$-URA3-$P_{T7}$-*aac(3)IV* | This work | MN450170 |
| pSEVA221 | $Km^R$; RK2 ori | 10 | JX560327 |
| pSEVA221T7RNAP | $Km^R$; pSEVA221, *tetR*-$P_{tetA}$-T7RNAP | This work | MN450171 |
| pSEVA221AID-T7RNAP | $Km^R$; pSEVA221, *tetR*-$P_{tetA}$-AID-T7RNAP | This work | MN450172 |
| pSEVA221pmCDA1-T7RNAP | $Km^R$; pSEVA221, *tetR*-$P_{tetA}$-pmCDA1-T7RNAP | This work | MN450173 |
| pSEVA221rAPOBEC1-T7RNAP | $Km^R$; pSEVA221, *tetR*-$P_{tetA}$-rAPOBEC1-T7RNAP | This work | MN450174 |
| pSEVA221TadA*-T7RNAP | $Km^R$; pSEVA221, *tetR*-$P_{tetA}$-TadA*-T7RNAP | This work | MN450175 |
| pdCas9 | $Cm^R$; p15A ori, *tracrRNA, cas9*(D10A, H840A), repeat-*Bsa*I spacer-repeat | 11 | Not deposited |
| pdCas9b.a | $Cm^R$; double spacer array Tb.a cloned in pdCas9 | This work | Not deposited |
| pdCas9b.a.c | $Cm^R$; triple spacer array Tb.a.c cloned in pdCas9 | This work | Not deposited |

#### Supplementary Table 3. Oligonucleotides to generate the spacer arrays

| Number | Name | Sequence (5’-3’) |
|---|---|---|
| **1** | F_bDRa | AAACAAATGAATTTCAGGGTCAGTTTGCCGTACGGTTTTAGAGCTATGCTGTTTTGAATGGTCCCAAAACATAACGTCACCGTCCAGTTCCACCAGAATG |
| **2** | R_bDRa | AAAACATTCTGGTGGAACTGGACGGTGACGTTAATGTTTTGGGACCATTCAAAACAGCATAGCTCTAAAACCGTACGGCAAACTGACCCTGAAATTCATTT |
| **3** | F_bDR | AAACAAATGAATTTCAGGGTCAGTTTGCCGTACGGTTTTAGAGCTATGCTGTTTTGAATGGTCCCAAAAC**GCTG** |
| **4** | R_bDR | GTTTTGGGACCATTCAAAACAGCATAGCTCTAAAACCGTACGGCAAACTGACCCTGAAATTCATTT |
| **5** | F_a*DR | ACGTCACCGTCCAGTTCCACCAGAATGTTTTAGAGCTATGCTGTTTTGAATGGTCCCAAAAC |
| **6** | R_a*DR | **ACTC**GTTTTGGGACCATTCAAAACAGCATAGCTCTAAAACATTCTGGTGGAACTGGACGGTGACGT**CAGC** |
| **7** | F_c* | **GAGT**GATTGCGTGCTCAGGTAATGATTGTCG |
| **8** | R_c* | AAAACGACAATCATTACCTGAGCACGCAATC |

Spacer Tb is shadowed in green, Ta (oligonucleotide 1 and 2) and Ta* (oligonucleotide 5 and 6) in cyan, and Tc* in yellow. DR nucleotides are underlined and the connector sequences are in red.

#### Supplementary Table 4. Spacers used in the arrays

| Name | Sequence (5’-3’) |
|---|---|
| Ta | ATTAACGTCACCGTCCAGTTCCACCAGAAT |
| Tb | AAATGAATTTCAGGGTCAGTTTGCCGTACG |
| Tc | CGCGGATTGCGTGCTCAGGTAATGATTGTC |
| Ta* | GCTGACGTCACCGTCCAGTTCCACCAGAAT |
| Tc* | GAGTGATTGCGTGCTCAGGTAATGATTGTC |

The connector sequences for assembly are underlined

**Supplementary Table 5. Oligonucleotides used in the allelic replacement method**

| Name | Sequence (5'-3') | Use |
|---|---|---|
| rph | A*G*AGCTACTCATCTTGTTGGCTCTGGCCCGAGGGG**G**AATCGAATCCATTGTAGCGACGCAGAAGGCGG*C*G | G insertion in *rph* |
| F_rph_A | CGCTACAATGGATTCGATTCCCCT | Allelic-specific PCR |
| F_rph_B | GCTACAATGGATTCGATTCCCCC | Allelic-specific PCR |
| R_rph | GTCGGAATTGTGAACGGCGAAG | Allelic-specific PCR |
| F_seq_pyrE | GCCTAACAGTGCCAGATCGCG | Amplification and sequencing of *pyrE* |

(*) Asterisks indicate phosphorothioate bonds.

**Supplementary Table 6. Oligonucleotides used for Sanger DNA sequencing.**

| Name | Sequence (5'-3') |
|---|---|
| F_GFPseq | GTACGTGGCGTCACCTTCACCC |
| R_T0ter | TACGAAGCTTCTGGATTCTCACCAATAAAAAACGCC |
| F_CS1_URA3 | ACACTGACGACATGGTTCTACAGGTGTGGTGGGTCCAGGTAT |
| R_CS2_URA3 | TACGGTAGCAGAGACTTGGTCTCGGCGTCATAATCAGCCAAT |

The Illumina tag sequences CS1 and CS2 are underlined.

#### Supplementary Figure 1

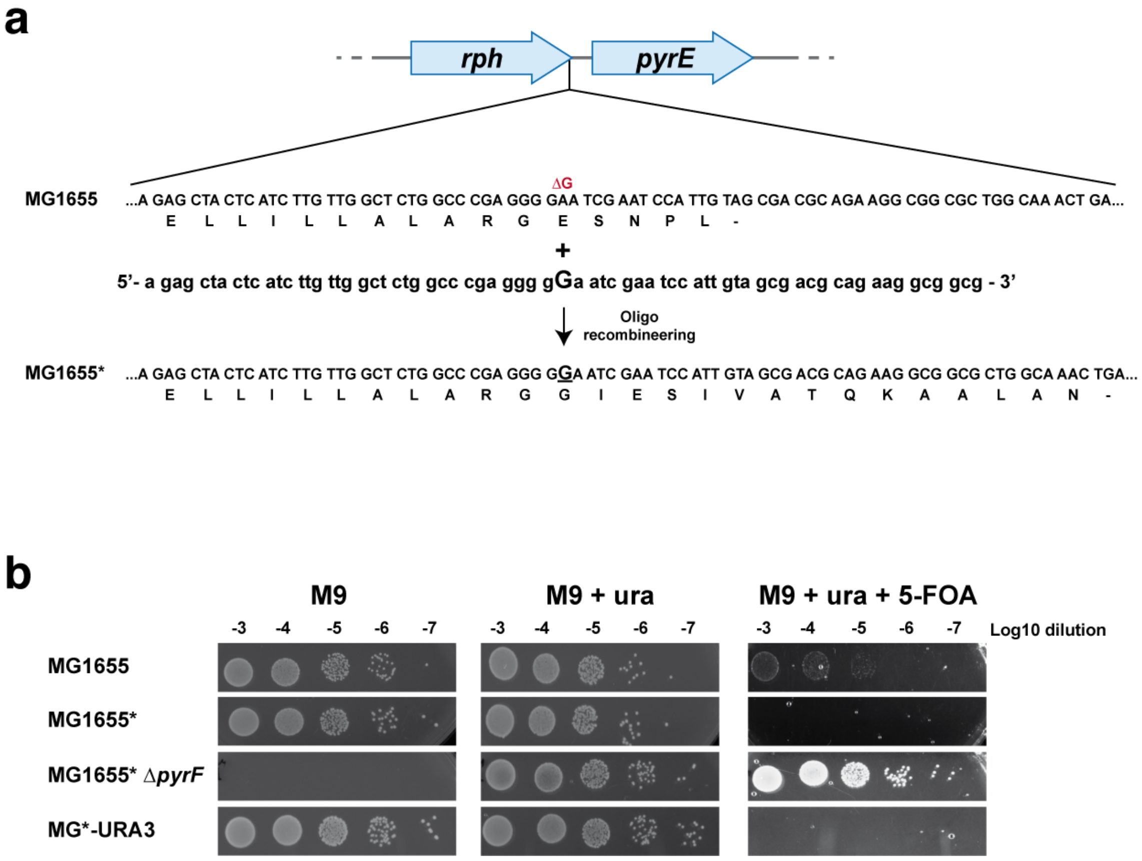


**Supplementary Figure 1.** Generation and characterization of the reporter strain MG*-URA3. (**a**) Scheme of the operon with the genes *rph* and *pyrE* from MG1655. It is shown the DNA sequence at the 5' end of *rph* with its amino acid translation below and the oligonucleotide used for G insertion resulting in the strain MG1655*. (**b**) Viability of different strains in minimal medium M9, M9 supplemented with uracil and M9 supplemented with uracil and 5-FOA. Series of ten-fold dilutions of each culture were prepared with 1X PBS and 10 μl drops of each dilution were plated.

**Supplementary Figure 2**

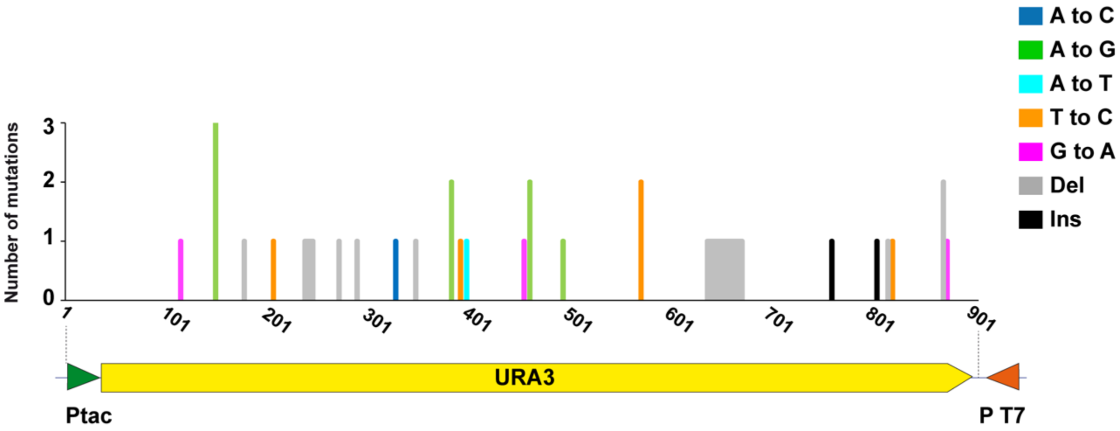


**Supplementary Figure 2**. Characterization of URA3 mutations in 30 FOA$^R$ colonies from strain MG*-URA3Δ*ung* expressing native T7RNAP. The position of the tac and T7 promoters are indicated (arrow heads). The identified mutations (following the color code on the right) are indicated with respect to the coding strand of URA3. Del: deletion; Ins: insertion.

**Supplementary Figure 3**

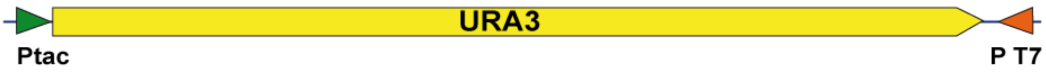


| | | |
|---|---|---|
| Coding strand of URA3<br>Non-coding strand of URA3 | 5’-GATC-3’<br>3’-CTAG-5’ | Template strand for T7RNAP<br>Non-template strand for T7RNAP |

↓

| | | |
|---|---|---|
| CD-T7RNAP<br>C to T in template strand: | 5’-GATT-3’<br>3’-CTAA-5’ | **C to T** in URA3 coding strand |
| CD-T7RNAP<br>C to T in non-template strand: | 5’-AATC-3’<br>3’-TTAG-5’ | **G to A** in URA3 coding strand |
| AD-T7RNAP<br>A to G in template strand: | 5’-GGTC-3’<br>3’-CCAG-5’ | **A to G** in URA3 coding strand |
| AD-T7RNAP<br>A to G in non-template strand: | 5’-GACC-3’<br>3’-CTGG-5’ | **T to C** in URA3 coding strand |

**Supplementary Figure 3**. Orientation of the DNA strands of URA3 in relation to the tac and T7 promoters.

**Supplementary Figure S4**

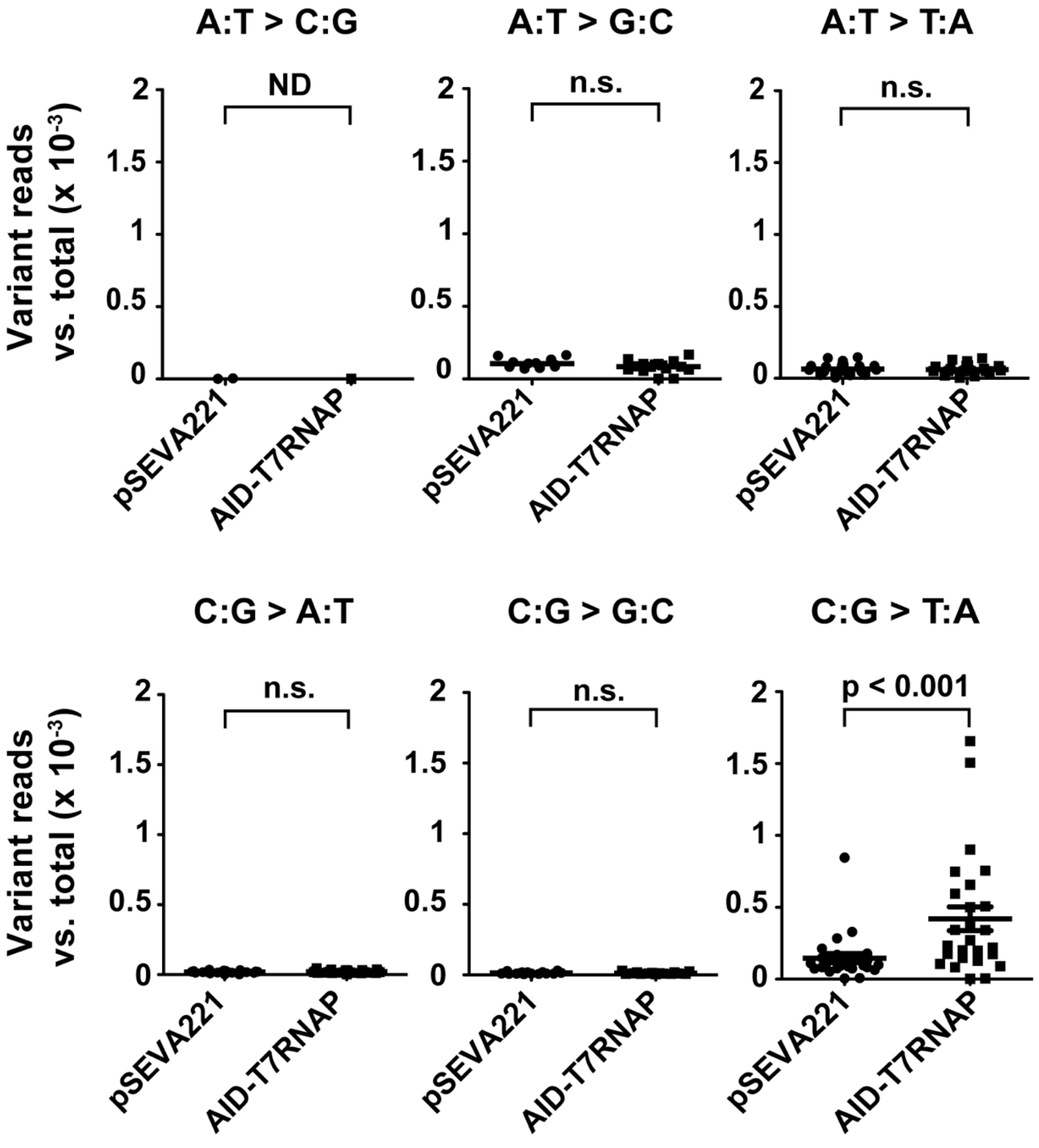


**Supplementary Figure 4.** Variant calling analysis of a 200 bp region of URA3 after its massive DNA sequencing (ca. $10^6$ reads) upon amplification from the strain MG*-URA3Δ*ung* with the empty plasmid (pSEVA221) or expressing AID-T7RNAP. The number of reads with different variants *vs*. total reads are represented with circles (empty plasmid) and squares (AID-T7RNAP). The lines represent the means and the standard errors from each group. The statistical analysis was done using the Mann Whitney test indicating p-value or n.s. (not significant). ND, not determined for groups with less than three variants detected.

**Supplementary Figure 5**

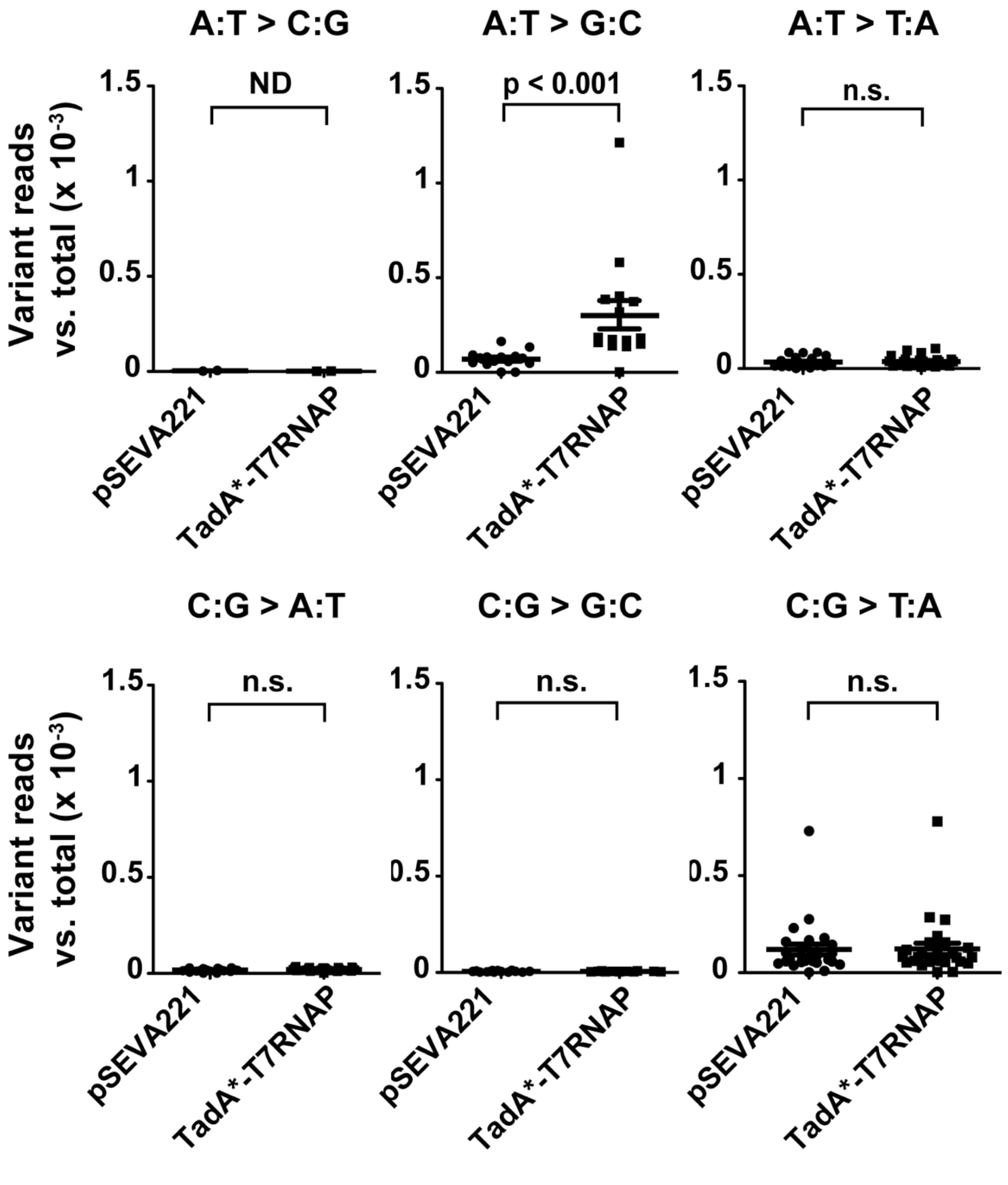


**Supplementary Figure 5**. Variant calling analysis of a 200 bp region of URA3 after its massive DNA sequencing (ca. $10^6$ reads) upon amplification from the strain MG*-URA3 $\Delta$*ung* $\Delta$*nfi* with the empty plasmid (pSEVA221) or expressing TadA*-T7RNAP. The number of reads with different variants *vs*. total reads are represented with circles (empty plasmid) and squares (TadA*-T7RNAP). The lines represent the means and the standard errors from each group. The statistical analysis was done using the Mann Whitney test indicating p-value or n.s. (not significant). ND, not determined for groups with less than three variants detected.

**Supplementary Figure 6**

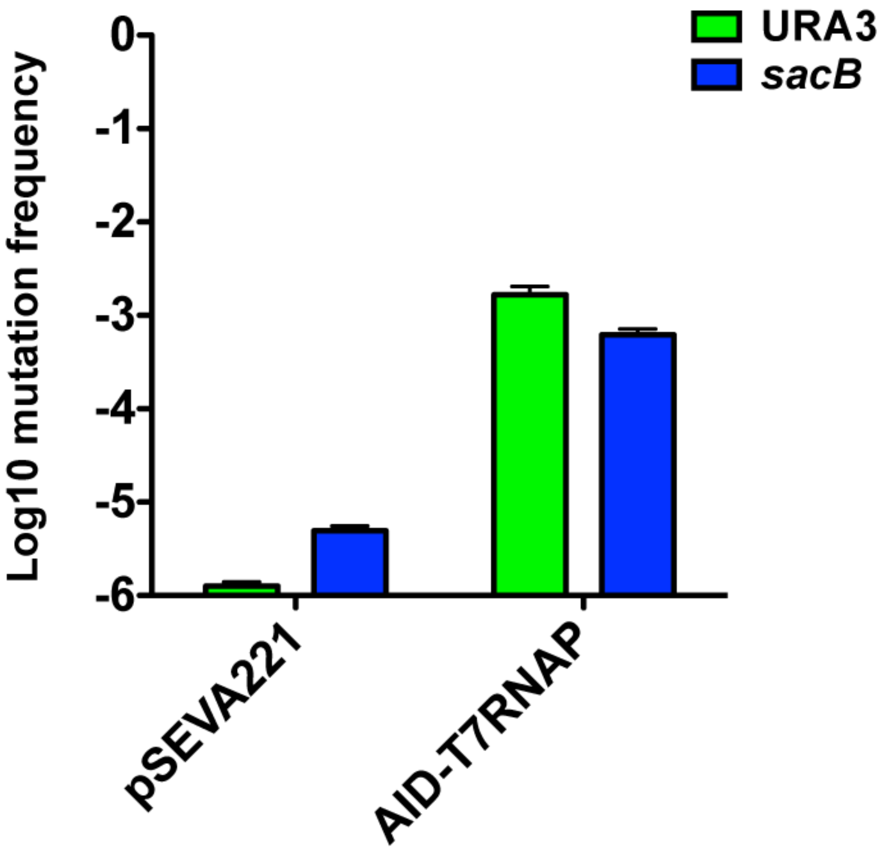


**Supplementary Figure 6.** Mutagenesis frequency in URA3 and *sacB* of the strain MG*-SacB-URA3Δ*ung*Δ*nfi* with pSEVA221 or expressing AID-T7RNAP.

#### Supplementary Figure 7

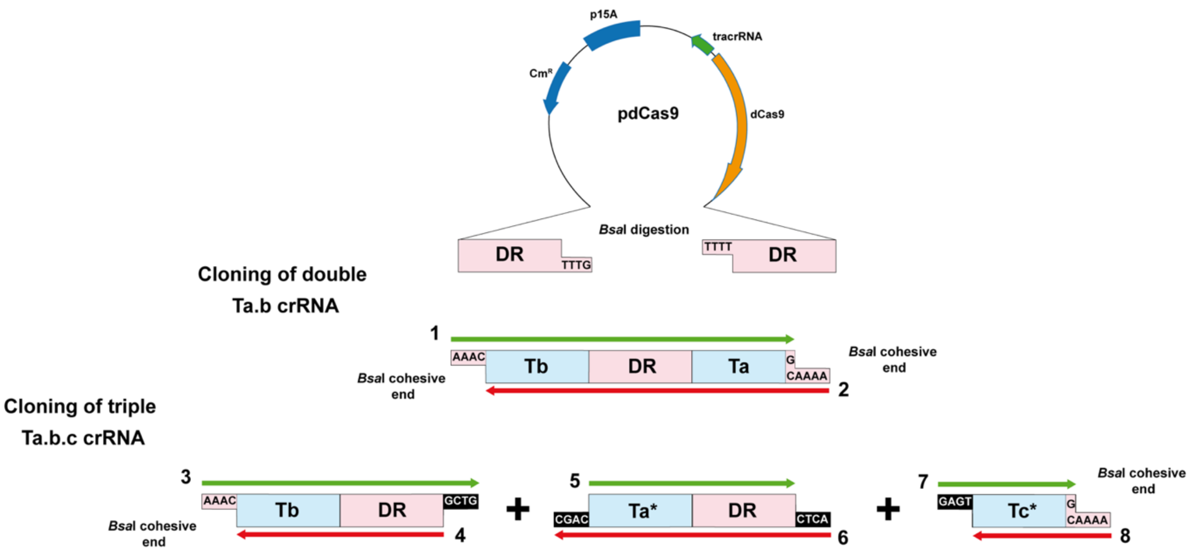


**Supplementary Figure 7.** Construction of pdCas9b.a and pdCas9b.a.c. Pink blocks represent direct repeats and blue blocks spacers. The complementary oligonucleotides are indicated with green and red arrows.

#### Supplementary Figure 8

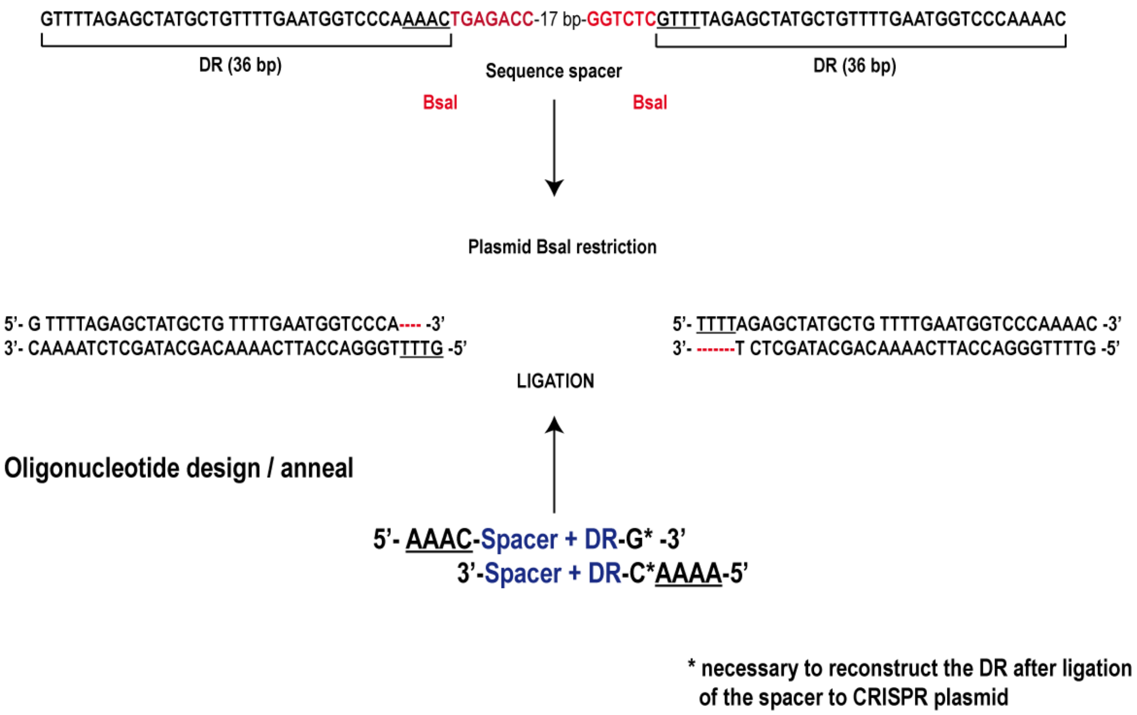


**Supplementary Figure 8.** Cloning the double and triple spacer arrays into the plasmid pdCas9 digested with *Bsa*I.
